# Biophysical modeling of actin-mediated structural plasticity reveals mechanical adaptation in dendritic spines

**DOI:** 10.1101/2022.11.06.515372

**Authors:** M. Bonilla-Quintana, P. Rangamani

**Author notes:** To whom correspondence must be addressed.

## Abstract

Synaptic plasticity is important for learning and memory formation; it describes the strengthening or weakening of connections between synapses. The postsynaptic part of excitatory synapses resides in dendritic spines, which are small protrusions on the dendrites. One of the key features of synaptic plasticity is its correlation with the size of these spines. A long-lasting synaptic strength increase (long-term potentiation, LTP) is only possible through the reconfiguration of the actin spine cytoskeleton. Here, we develop an experimentally-informed three-dimensional computational model in a moving boundary framework to investigate this reconfiguration. Our model describes the reactions between actin and actin-binding proteins (ABPs) leading to the cytoskeleton remodeling and their effect on the spine membrane shape to examine the spine enlargement upon LTP. Moreover, we find that the incorporation of perisynaptic elements enhances spine enlargement upon LTP, exhibiting the importance of accounting for these elements when studying structural LTP. Our model shows adaptation to repeated stimuli resulting from the interactions between spine proteins and mechanical forces.

**Significance Statement:** Dendritic spines are small protrusions that receive stimulation from presynaptic neurons. Upon stimulation, the dendritic spines change their size, an important feature of synaptic plasticity. This change is achieved by modifications to the actin cytoskeleton and mediated by many actin-binding proteins. To investigate the fundamental mechanics of spine expansion, we developed a 3D biophysical model that accounts for the dynamics of cytoskeleton-membrane interactions. Our simulations predict that spine expansion due to actin remodeling can be enhanced by including the interaction with perisynaptic elements that affect the spine’s mechanical properties. We also found that mechanical properties can control spine expansion after repeated stimuli, which ensures physiological size. Thus, we predict that spine growth is regulated by its mechanical properties.

## 1 Introduction

Dendritic spines are small protrusions from dendrites that form the postsynaptic part of a vast majority of excitatory synapses (Harris 2020; Nakahata and Yasuda 2018; Okabe 2020). In response to glutamate signals released by stimulated presynaptic neurons, spines undergo both biochemical and morphological changes that can be long-lasting (Matsuzaki et al. 2004; Okamoto et al. 2004; Kasai et al. 2010) (Fig. 1). Such changes have been long hypothesized as the biological mechanisms underlying memory storage in the brain (Yuste and Bonhoeffer 2001). One of the most-studied long-lasting changes is long-term potentiation (LTP).

**Figure 1:**
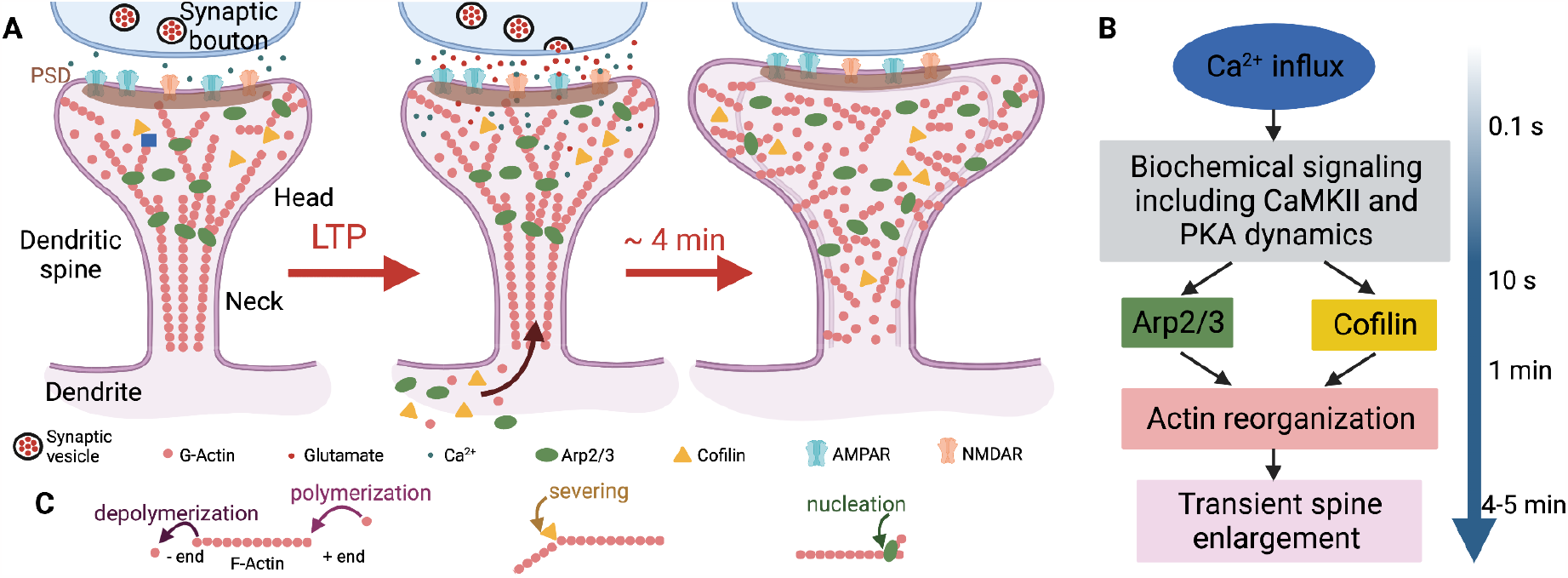
Biophysical events involving early structural plasticity in the dendritic spine. **A)** Upon LTP induction, glutamate is released from the presynaptic neuron and taken up by the postsynaptic neuron. Consequently, a cascade of chemical reactions is initiated **(B)**, and an influx of actin, cofilin, and Arp2/3 into the spine is triggered. This remodels the spine cytoskeleton allowing for spine enlargement. **C)** Actin filament treadmilling, branching, and severing events remodel the cytoskeleton. In this model, we focus on how the spatio-temporal dynamics of actin, cofilin, and Arp2/3 dictate the shape of the dendritic spine. Image created with BioRender.com

LTP induction alters the dendritic spine morphology. It has been shown that spines increase their volume dramatically, by up to 390%, after stimulation (Tønnesen et al. 2014; Chang et al. 2017). This increase is associated with an increase in AMPAR-mediated currents and depends on NMDAR activation and actin polymerization (Matsuzaki et al. 2004). Actin is highly concentrated in dendritic spines (Matus et al. 1982) and plays a key role in LTP (Cingolani and Goda 2008; Okabe 2020). Around 95% of F-actin in the spine undergoes rapid treadmilling (time scale ∼40 s), generating an expansive force caused by actin polymerization (Honkura et al. 2008). Moreover, the equilibrium between actin monomers (G-actin) and F-actin is affected by stimulation (Okamoto et al. 2004), suggesting that structural reconfiguration of actin is necessary for spine enlargement.

Actin-binding proteins (ABPs) aid the reconfiguration of the cytoskeleton by promoting G-actin polymerization, F-actin depolymerization, and capping the F-actin ends, which prevents their polymerization and depolymerization (Pollard and Borisy 2003). Experiments have shown that ABPs are necessary for LTP (Cingolani and Goda 2008; Fortin, Srivastava, and Soderling 2012). To achieve the transient size increase seen during the first 4-7 minutes of LTP (Tønnesen et al. 2014; Chang et al. 2017) translocation of ABPs into the spine is necessary (Bosch et al. 2014), in addition to the Arp2/3 and cofilin activation due to Ca^2+^ influx (Rangamani, Levy, et al. 2016) (Fig. 1C). After three hours, the spine settles to a size that is 40% larger than that prior to LTP induction (Tønnesen et al. 2014; Chang et al. 2017). Hence, the reconfiguration of the actin cytoskeleton of dendritic spines, and thus, structural plasticity is possible by an orchestrated interplay between actin and ABPs triggered by different signaling pathways (Bosch et al. 2014).

Besides changes in size promoted by the cytoskeleton reconfiguration, spines experience further mechanical modifications upon LTP. For example, mechanical coupling of the actin filaments with the extracellular environment through molecular clutches is necessary to push the membrane forward (Kastian et al. 2021). Degradation of the extracellular matrix (ECM) by proteases promotes structural and functional LTP (Wang et al. 2008). The elastic storage modulus and viscous loss modulus of the spine increase, which facilitates its mechanical stabilization due to the stiffening of the internal structure (Smith et al. 2007). Therefore, both biochemical reactions and mechanical force generation are required for structural LTP (sLTP).

In this work, we seek to answer the following questions: Can a minimal model of actin-membrane interactions capture the dynamics of spine changes during sLTP? How spine enlargement upon LTP can be enhanced by the mechanical changes induced through the interaction with other perisynaptic elements? And finally, how do repeated stimuli affect sLTP? To answer these questions, we develop a 3D computational model using a system of partial differential equations with moving boundaries to incorporate the spatio-temporal dynamics of actin and ABPs in the expanding dendritic spines. We systematically investigate the contribution of ABPs and mechanics to sLTP under different conditions that mimic the alteration of spine properties. Our results predict that actin interaction with ABPs is sufficient to capture the spine growth during sLTP and that further increase can be obtained when including the interaction with other perisynaptic elements. Moreover, the spine enlargement capacity diminished with repeated stimuli, hinting at a homeostatic mechanism related to its mechanical properties.

## 2 Materials and Methods

We develop a mathematical model in which F-actin dynamics in the dendritic spine are affected by Arp2/3 and cofilin. As in cell motility models (Mogilner and Edelstein-Keshet 2002; Tania, Prosk, et al. 2011; Tania, Condeelis, and Edelstein-Keshet 2013), we assume that these ABPs are sufficient to promote membrane protrusion because their interaction with F-actin increases the force generated by actin polymerization which helps to overcome the membrane resistance (Xiong et al.2010; Mogilner and Oster 1996). Thus, we expect that the spine enlargement seen shortly after LTP induction will be driven by the reconfiguration of the cytoskeleton, similar to the reconfiguration needed for cell motility, and that other proteins minimally affect spine expansion. In the model, actin, Arp2/3, and cofilin are free to diffuse in the spine volume. Upon LTP, there is an influx of these proteins into the spine that represents the translocation of ABPs. Note that the ABPs chosen in our model are required for synaptic function. For example, conditional mutagenesis of Arp2/3, which promotes actin branching (Pollard and Borisy 2003; Pollard, Blanchoin, and Mullins 2000), hinders spine enlargement upon LTP and is implicated in psychiatric disorders (Kim, Racz, et al. 2013). Loss of cofilin impairs learning (Rust et al. 2010). Cofilin is an ABP whose function depends on its relative concentration with actin; it severs F-actin at low concentrations but promotes actin nucleation at high concentrations (Andrianantoandro and Pollard 2006).

### 2.1 Governing equations

Based on the assumptions described above, we formulate a system of partial differential equations (PDEs) that describe the spatio-temporal dynamics of F-actin with uncapped (+) ends (or barbed ends), Arp2/3, and cofilin. Note that barbed ends polymerize G-actin, which generates an expanding force. Moreover, the action of Arp2/3 and cofilin, in addition to the actin influx upon LTP induction, increase the number of barbed ends in the spine. Thus, in our model, the increment of barbed ends during stimulation allows us to approximate the dynamics of barbed ends with PDEs instead of considering single filaments. This system is coupled with the spine membrane dynamics, as described in (Doubrovinski and Kruse 2011). To examine the dendritic spine size and shape changes, we implement a moving boundary framework. The forces generated by actin polymerization **F**_*actin*_, the membrane **F**_*mem*_, and drag **F**_*drag*_ dictate the displacement of the membrane Γ, which enclose the spine to an evolving domain Ω, hence Γ = *∂*Ω.

In the absence of inertia, the force balance becomes

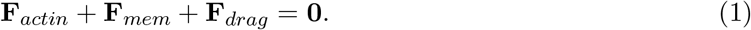

Note that vectors are in bold font. As in (Gonçalves and Garcia-Aznar 2021), we describe the interactions between the spine and the extracellular environment through **F**_*drag*_, a dissipative force that can represent the contributions of fluid drag and extracellular matrix adhesion. The drag force is given by

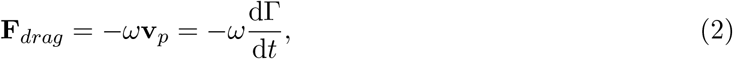

where *ω* represents an effective drag coefficient and **v**_*p*_ is the protrusion velocity, i.e., the displacement of the membrane over time. Thus, the membrane evolves according to

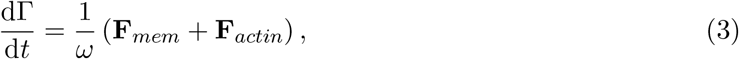

where **F**_*actin*_ is the force generated by the polymerization of F-actin near the dendritic spine membrane that pushes the membrane forward (Mogilner and Oster 1996; Lacayo et al. 2007; Honkura et al. 2008), given by

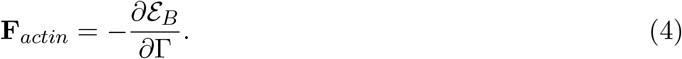

Following (Doubrovinski and Kruse 2011), the force applied to the membrane is dictated by the total potential generated by the number of barbed ends per cubic volume *B* inside the spine

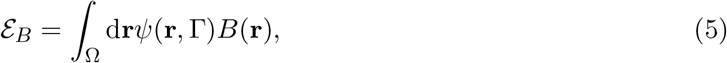

where

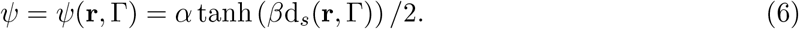

is a soft repulsive potential (Fig. 2L) that depends on the distance d_*s*_(**r**, Γ) (Fig. 2B-C), given by

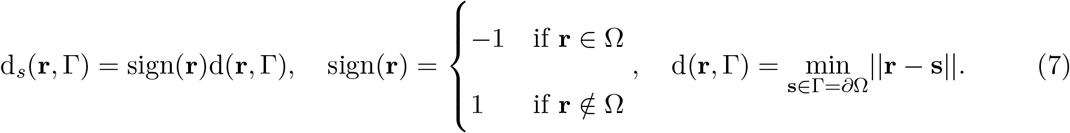

Thus, barbed ends closer to the membrane have greater contributions to the potential energy. Note that when *α, β* 2212→ ∞, the membrane is a reflective boundary. We use a signed distance to track the relative position of **r** with respect to the spine membrane.

**Figure 2:**
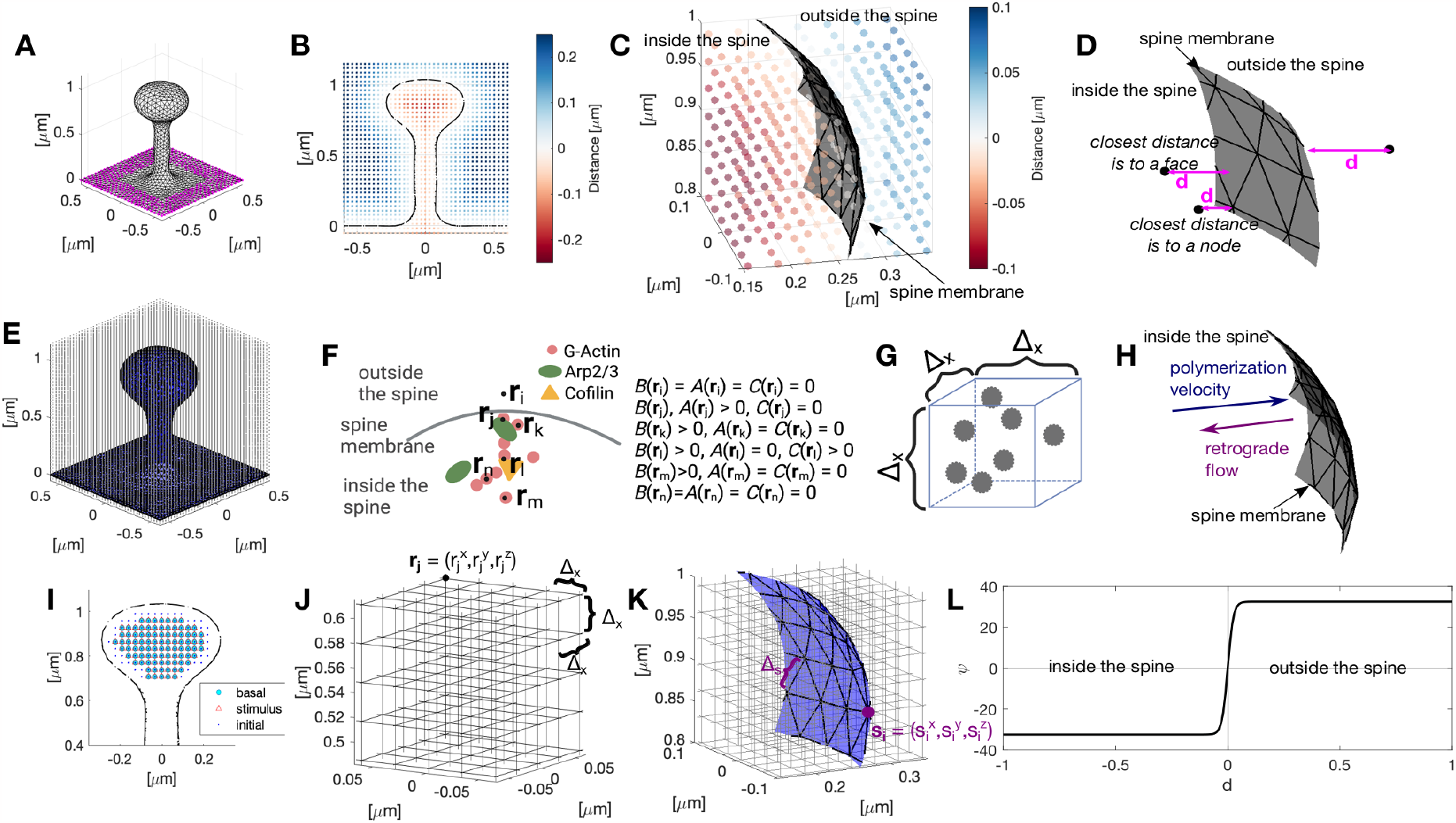
Simulation setup. **A)** Initial configuration of the dendritic spine. Magenta dots correspond to the nodes of the dendritic shaft that remain fixed throughout the simulation. **B**) Slide of a dendritic spine at x = 0 µm showing the value of the signed distance function in the cubic domain ℒ . **C**) Zoom of B. Each dot represents a position r, color-coded to the signed distance from the membrane (gray mesh). **D**) Scheme of the different cases for calculating the distance to the spine membrane. Arrows correspond to d =∇ds. **E**) Dendritic spine (blue mesh) embedded in a cubic domain ℒ. Dots correspond to the positions r (the discretization of this domain). **F**) We assume a given concentration of Arp2/3, cofilin and number of barbed ends per discretized volume (**G**), instead of modeling the individual filaments. H) Scheme of the different types of motion for F-actin. **I**) Slide of a dendritic spine at x = 0 µm showing the spatial locations of basal influx (cyan circles) and stimulus-triggered influx (red triangles). Blue dots indicate the initial position of protein densities. Note that the red triangles, blue dots, and cyan circles overlap and that the stimulus-triggered influx is homogeneous in the spine head. **J**) Zoom to the cubic domain in ℒ E, discretized in intervals of length Δ _x_. The intersection of the grid lines corresponds to the discretized positions **r. K**) Zoom of the spine membrane (in blue) embedded in the cubic domain. This membrane is discretized using a triangular mesh, where the node positions s are time evolving. **L**) Plot of the function ψ in Eq. (6).

The force generated by actin is balanced by an opposing force generated by the membrane **F**_*mem*_, which counteracts membrane deformations and is given by

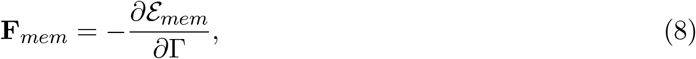

where

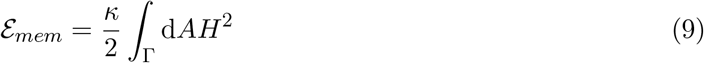

represents the membrane Helfrich free energy due to bending (Helfrich 1973). Here, *H* is the mean curvature, and *κ* is the bending modulus. Note that we consider the membrane as a 2D elastic continuum with negligible thickness, as in (Krüger 2012; Deserno 2015). The volume and surface area of the spine membrane are unconstrained due to the influx of proteins and membrane addition through trafficking mechanisms (Yang and Liu 2022). Thus, we do not consider membrane tension and osmotic pressure in our model.

#### 2.1.1 Actin dynamics

F-actin in dendritic spines is distributed in two different pools, the stable pool and the dynamic pool. In contrast to the stable pool of actin, with a lifetime of ∼17 min, the dynamic pool undergoes rapid treadmilling (∼40 s) (Honkura et al. 2008). We assume that the structural changes in dendritic spines are mostly driven by the dynamic pool that has short filaments with uncapped (+) ends undergoing continuous polymerization, which generate **F**_*actin*_. Therefore, we do not explicitly model the stable pool, which only accounts for 5% of the total F-actin (Honkura et al. 2008). We keep track of the number of barbed ends per unit volume *B*(**r**, *t*) at position **r** ∈ L ⊂ ℝ^3^, where L is a cubic lattice domain (see Fig. 2F-G), instead of F-actin or G-actin concentration. Note that keeping track of the number of barbed ends instead of the number of actin filaments reduces the complexity of the model because otherwise, we would have to account for the length of the filaments. Here, we assume that actin filaments with barbed ends have similar lengths. Although this is a simplifying assumption, the modeled F-actin is dynamic and has a short length (Honkura et al. 2008), which we expect to exert similar force to the membrane; hence, differences in F-actin length are negligible. Lastly, we assume that ATP is not depleted throughout the simulation.

The dynamics of the number of barbed ends per unit volume are given by

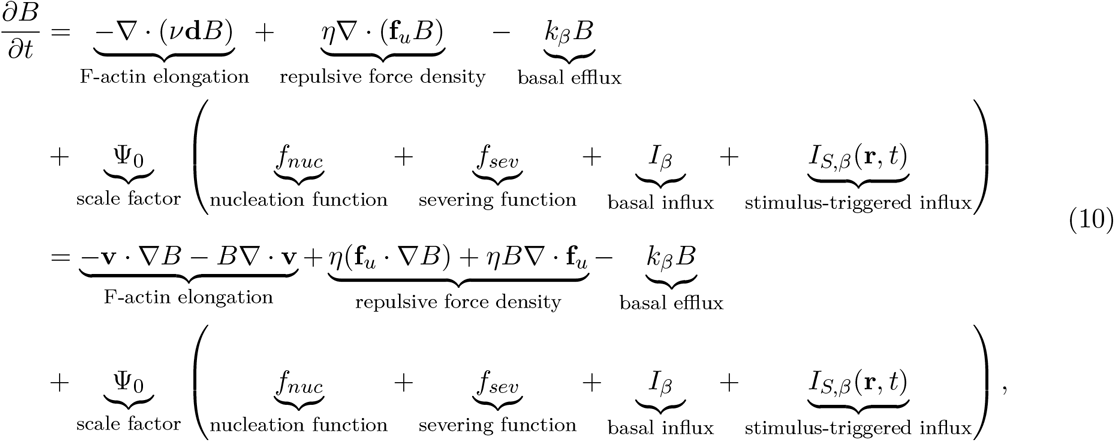

where *k*_*β*_ is a constant basal efflux rate. The basal influx *I*_*β*_ of actin is due to local actin synthesis (Cajigas et al. 2012; Tiruchinapalli et al. 2003). Because the proteins used in the model are dispersed in the spine head, we assume a homogeneous basal influx. The stimulus-triggered influx *I*_*S,β*_ mimics the transient influx upon stimulation (Bosch et al. 2014). Hence, it is nonzero only during a brief window of time after stimulus initiation (1 minute). In line with experimental observations (Bosch et al. 2014), we localize the stimulus-triggered influx to the spine head. For consistency of units, these terms are multiplied by a conversion factor Ψ_0_ that changes from concentration units in µM to the number of barbed ends per µm^3^. ∇ represents the gradient operator. Note that the number of barbed ends per unit volume is not conserved due to the stimulus-triggered influx.

The first term in Eq. (10), 2212∇ · (*ν***d***B*), denotes the change in the barbed ends per unit volume due to the polymerization of G-actin in the direction **d** = **d**(**r**, Γ) = ∇d_*s*_(**r**, Γ), d_*s*_ is defined in Eq. 7 (Fig. 2D). Hence, **d** is the unit vector emanating from **r** and directed to the closest point in the membrane. F-actin (+) ends are continuously polymerizing G-actin at a speed *ν*. Since the dynamic pool treadmills fast and accounts for 95% of F-actin in the spine (Honkura et al. 2008), we take the actin polymerization velocity to be fixed and independent of actin concentration. Thus, **v** = *ν***d** represents the velocity field of actin polymerization.

The second term in Eq. (10), *η*∇ · (**f**_*u*_*B*), accounts for the change in *B* due to a force density **f**_*u*_, given by

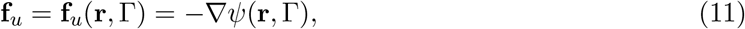

where *ψ* is the is a soft repulsive potential defined in Eq. (6). Actin filaments interact with the intracellular environment, which includes transient attachments to the substrate (Doubrovinski and Kruse 2011). The effects of such interactions are represented by effective filament motility parameter *η* that reduces the impact of ∇ · **f**_*u*_ on *B*. The force field **f**_*u*_ confines the system to Ω (Doubrovinski and Kruse 2011). Note that the vectors of **f**_*u*_ have direction **d** (Fig. 2D,H). Hence, **f**_*u*_ can describe the force that generates a retrograde flow of F-actin to create a gap between the barbed end and the membrane to fit G-actin during polymerization.

The nucleation function accounts for the nucleation of new filaments with uncapped (+) ends by Arp2/3 binding to actin filaments. We choose the nucleation function proposed by (Carlsson, Wear, and Cooper 2004)

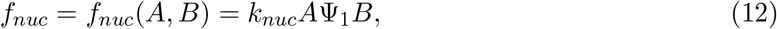

which reflects the side-branching of Arp2/3. This function agrees with the observations in (Risca et al. 2012), where the nucleation of new filaments is enhanced on the side of bent filaments. Here, *k*_*nuc*_ represents the nucleation rate, *A* is the Arp2/3 concentration, and Ψ_1_ is a unit conversion factor.

Cofilin severs F-actin in a concentration-dependent manner (Andrianantoandro and Pollard 2006), creating new filaments with uncapped (+) ends. Based on observations of (Bosch et al. 2014), we assume that during the first few minutes after LTP the concentration of F-actin in the spine is higher than the concentration of cofilin. Because cofilin severs F-actin at low concentrations (Andrianantoandro and Pollard 2006), we assume that cofilin binding to F-actin is cooperative (De La Cruz 2005). This is represented by the severing rate function

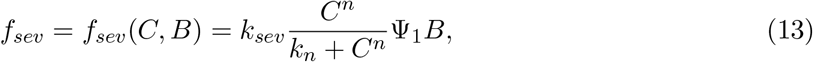

where *k*_*sev*_ is the severing rate, *k*_*n*_ is the dissociation constant, *n* is the Hill coefficient to capture the cooperative nature of the kinetics, and *C* represents the concentration of cofilin.

#### 2.1.2 Arp2/3 Dynamics

The evolution of Arp2/3 concentration over time is given by

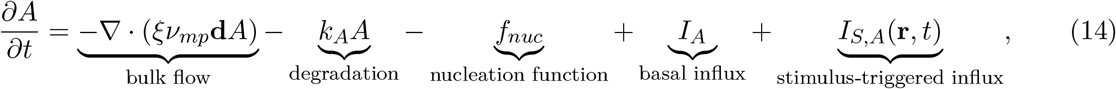

where *k*_*A*_ is a constant degradation rate. We assume that as the dendritic spine expands, the Arp2/3 molecules are transported towards the membrane by a bulk flow, as in (Tania, Condeelis, and Edelstein-Keshet 2013). Note that the motion of the bulk flow is driven by the motion of F-actin since actin is highly dense in the spine. The speed of the bulk flow is given by *ξν*_*mp*_, with *ν*_*mp*_ representing the protrusion velocity. To ease the model simulations, we make the following simplifications: 1) instead of calculating *ν*_*mp*_ at each node of the mesh representing the spine membrane, we take the velocity of the protrusion at one side of the spine head for all nodes, as described in Section 2.4. 2) If the protrusion shrinks at that location, i.e., the velocity direction is opposite to the expanding direction, we take *ν*_*mp*_ = 0 to avoid numerical instability. 3) The polymerization velocity (*ν* in Eq. 10) is lower than *ν*_*mp*_. Thus, we multiply *ν*_*mp*_ by 0 *< ξ <* 1 to have similar velocities in all the variables and avoid numerical problems. This can represent the hindering of Arp2/3 and cofilin by the high density of proteins inside the dendritic spine (Helm et al. 2021).

#### 2.1.3 Cofilin Dynamics

The evolution of cofilin concentration is given by

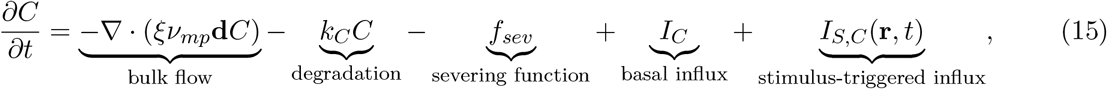

where *k*_*C*_ is the degradation rate. Note that cofilin is also transported by bulk flow towards the membrane at speed *ξν*_*mp*_.

### 2.2 Numerical implementation

In the model, the dynamics of actin barbed ends, Arp2/3, and cofilin are given by Eqs. (10),(14), and (15). In the simulation, these equations are solved over time in a cubic domain with an embedded triangular mesh representing the dendritic spine membrane (Fig. 2E). A signed distance function is used to calculate the proximity of the chemical species to the membrane (Fig. 2B). The force generated by actin polymerization and the force generated by the membrane are calculated from the spatial location of the barbed ends and the geometry of the spine mesh, respectively. These forces dictate the evolution of the spine mesh (Eq. 3). To reach a stable spine morphology, we set the basal influx to be homogeneously distributed inside the spine head and impose absorbing boundary conditions because the proteins have a stable arrangement in the spine neck (Bär et al. 2016). The basal influx remains in the same location throughout the simulation (cyan circles in Fig. 2I). We assume that the stimulus-triggered influx rapidly reaches the spine head at Δ_*x*_ distance from the membrane and within *z* = 0.7 and *z* = 1 µm (red triangles in Fig. 2I). We restrict the height of the stimulus location to capture the observations that the configuration of the postsynaptic density (PSD), a dense receptor site at the tip of the spine, remains stable during the early phase of sLTP (Bosch et al. 2014; Tønnesen et al. 2014), i.e., the duration of the simulations. Finally, we scale the stimulus-triggered influx to the initial head size to ensure that the same amount of proteins flows into the spine during the stimulus time window.

We solve the system of partial differential equations using MATLAB’s (MATLAB 2021) `ode45` solver at each time-step Δ_*t*_ for all positions **r** of a discretized cubic lattice domain L ∈ ℝ^3^ with Δ_*x*_ spacing (Fig. 2J). The gradient operator is discretized using an explicit finite difference scheme. The spine membrane Γ is approximated by a 3D polygon with a triangular isotropic mesh consisting of *n*_*v*_ vertices located at **s**_*i*_ ∈ ℝ^3^, *i* ∈ {1, 2, …, *n*_*v*_} (Fig. 2K). Γ is updated according to Eq.(3) at each time-step. For numerical accuracy, the membrane is remeshed using an isotropic remesher (Helf 2021) (based on OpenMesh OpenMesh 2020), with a target edge length of Δ_*s*_. The points corresponding to the base of the dendrite are fixed throughout the simulation (Fig. 2A). See Figure 3 for a flowchart of the simulation.

**Figure 3:**
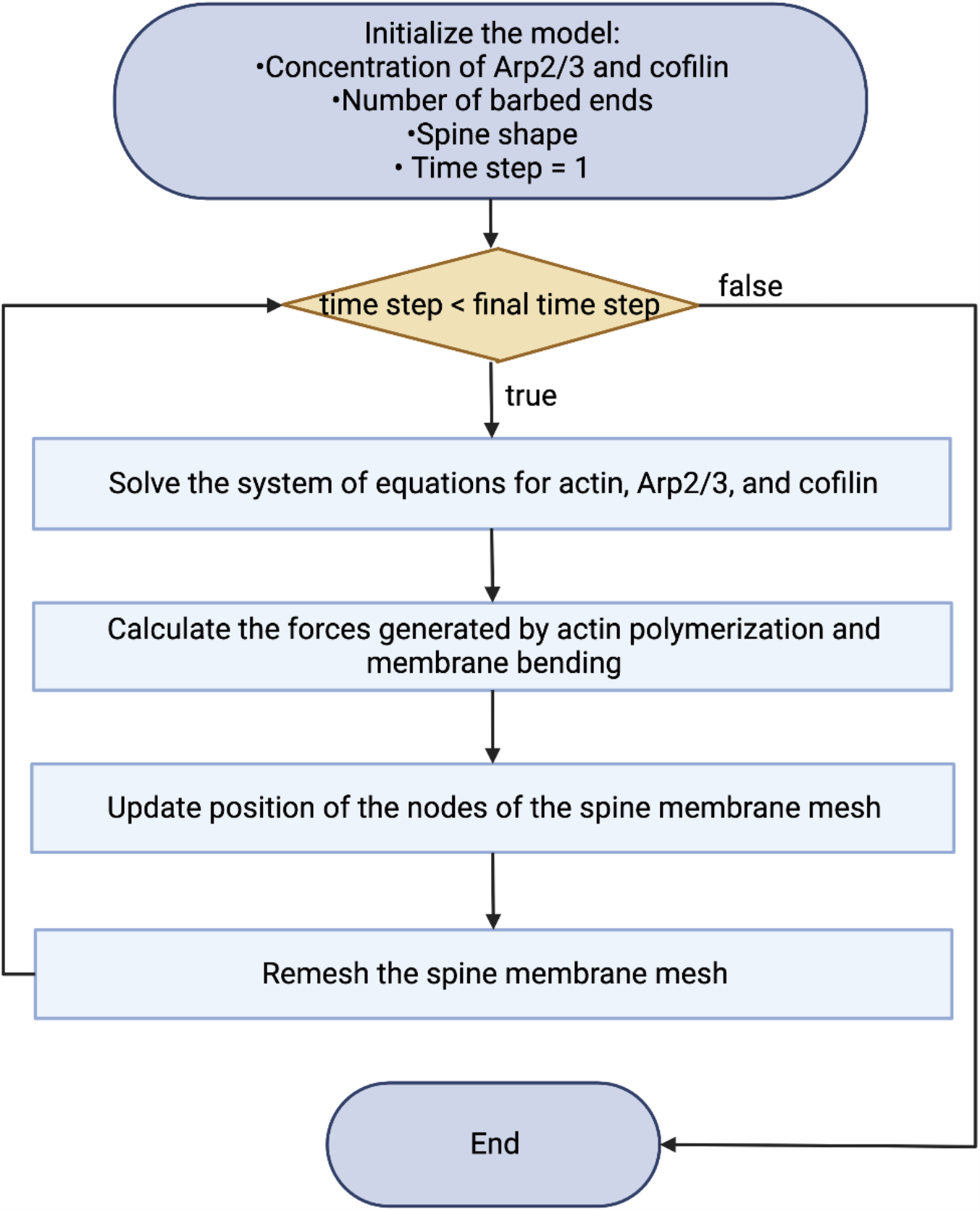
Flowchart outlining the key steps of the simulation. The full code is available at Zenodo (https://zenodo.org/records/10114856, version 2).

#### 2.2.1 Membrane Discretization

In this section, we present the discretized version of the continuous PDE system that we implement in the simulation. Note that we use the concentration of Apr2/3 and cofilin, or the number of barbed ends per unit volume (Fig. 2G) instead of modeling the structure of the dendritic spine cytoskeleton (Fig. 2F). The points of the cubic domain mesh **r** remain constant over the simulation (Fig. 2E,J). A triangular mesh that represents the spine membrane is embedded in this cubic domain (Fig. 2K). The position of the nodes of the spine mesh **s** change at each time-step according to Eq. (3).

The distance d(**r**, Γ), defined in Eq. (7), corresponds to the minimum distance from the lattice point **r** to the hexagonal mesh representing the membrane Γ. For its calculation, there are two cases (Fig. 2D):

1. The closest point to the membrane from **r** is a vertex **s**. Then,

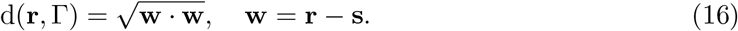

2. The closest point to the membrane from **r** is the triangular face *i* that spans through the vertices (**s**_*i*_, **s**_*i*+1_, **s**_*i*+2_). Then,

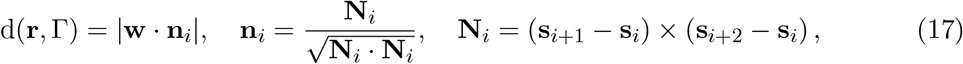

where **N**_*i*_ is the normal vector and × represents the cross product.

In our numerical implementation, the normal vector to the mesh surface points to the outside of the spine, and the polymerization direction is 2212**w** because we are using the signed distance. This is considered in the calculations by changing the signs accordingly.

For the calculation of the force generated by the membrane **F**_*mem*_ (Eq. 8), we follow (Zhu, Lee, and Rangamani 2022) and take the bending force with the spontaneous curvature equal to zero. To obtain the initial spine shape, we construct a mesh with a spheroid connected to the x-y plane representing the membrane via a cylinder, which represents the spine neck. To obtain a stable shape, we let the position of the vertices evolve according to Eq. (3), but only considering the force generated by the membrane (i.e., **F**_*actin*_ = **0**). We anchor the spine to the dendrite by fixing the vertices corresponding to the dendrite (magenta points in Fig. 2A).

#### 2.2.2 Code Accessibility

The code has been deposited in Zenodo (https://zenodo.org/records/10114856).

### 2.3 Constraining kinetic parameters to experimental measurements

Experimental results from (Bosch et al. 2014) form the foundation of the temporal dynamics of the number of barbed ends, and Arp2/3 and cofilin concentration in our model. Therefore, we fit the model parameters to the data in (Bosch et al. 2014) that shows the normalized concentrations of *β*-actin, Arp2/3, and cofilin-1 in dendritic spines of hippocampal CA1 neurons after inducing sLTP by 2 photon glutamate uncaging for 1 minute (Fig. 4A-B). Other parameters are taken from the literature or set to a physiological range (Table 1).

**Table 1:** Model Parameters. For further information about parameter fitting see Section 2.3.

| Symbol | Definition | Units | Value | Reference |
| --- | --- | --- | --- | --- |
| $k_\beta$ | actin degradation rate | 1/s | 0.0081 | fitted to (Bosch et al. 2014) |
| $I_\beta$ | actin basal influx | $\mu\text{M/s}$ | 24.4284 | fitted to (Bosch et al. 2014) |
| $I_{S,\beta}$ | actin stimulus influx | $\mu\text{M/s}$ | 25.6684 | fitted to (Bosch et al. 2014) |
| $k_A$ | Arp2/3 degradation rate | 1/s | 0.0013 | fitted to (Bosch et al. 2014) |
| $I_A$ | Arp2/3 basal influx | $\mu\text{M/s}$ | 0.0255 | fitted to (Bosch et al. 2014) |
| $I_{S,A}$ | Arp2/3 stimulus influx | $\mu\text{M/s}$ | 0.0293 | fitted to (Bosch et al. 2014) |
| $k_C$ | cofilin degradation rate | 1/s | 0.0006 | fitted to (Bosch et al. 2014) |
| $I_C$ | cofilin basal influx | $\mu\text{M/s}$ | 0.0237 | fitted to (Bosch et al. 2014) |
| $I_{S,C}$ | cofilin stimulus influx | $\mu\text{M/s}$ | 0.4384 | fitted to (Bosch et al. 2014) |
| $k_{nuc}$ | nucleation rate | $1/(\mu\text{M s})$ | 0.0153 | (Carlsson, Wear, and Cooper 2004) |
| $k_{sev}$ | cofilin severing rate | 1/s | 0.0120 | (Roland et al. 2008) |
| $n$ | Hill coefficient for cofilin binding | - | 3.5 | (De La Cruz 2005) |
| $k_n$ | dissociation constant | $\mu\text{M}^n$ | 0.6 | (De La Cruz 2005) |
| $\Psi_0$ | scale factor converting to number of barbed ends per $\mu\text{m}^3$ | $\#/(\mu\text{m}^3\mu\text{M})$ | 3.6 | fitted |
| $\Psi_1$ | scale factor converting to concentration | $\mu\text{m}^3\mu\text{M}/\#$ | 0.02 | fitted |
| $\nu$ | F-actin polymerization velocity | $\mu\text{m/s}$ | $1 \times 10^{-5}$ | fitted |
| $\xi$ | speed restriction | - | 0.1 | fitted |
| $\alpha$ | amplitude of the repulsive potential | pN | $3 \times 10^4$ | fitted |
| $\beta$ | steepness of the repulsive potential | - | 40 | fitted |
| $\eta$ | effective filament mobility | $\mu\text{m}/(\text{s pN})$ | $1 \times 10^{-10}$ | fitted |
| $\omega$ | drag coefficient | $\text{s pN} / \mu\text{m}$ | $1 \times 10^5$ | fitted |
| $\kappa$ | bending modulus | pN $\mu\text{m}$ | 0.18 | (Pontes et al. 2013) |
| $\Delta_t$ | time-step | s | 0.01 | fitted |
| $\Delta_x$ | cubic lattice interval length | $\mu\text{m}$ | 0.0315 | fitted |
| $\Delta_s$ | edge length | $\mu\text{m}$ | 0.05 | fitted |

**Figure 4:**
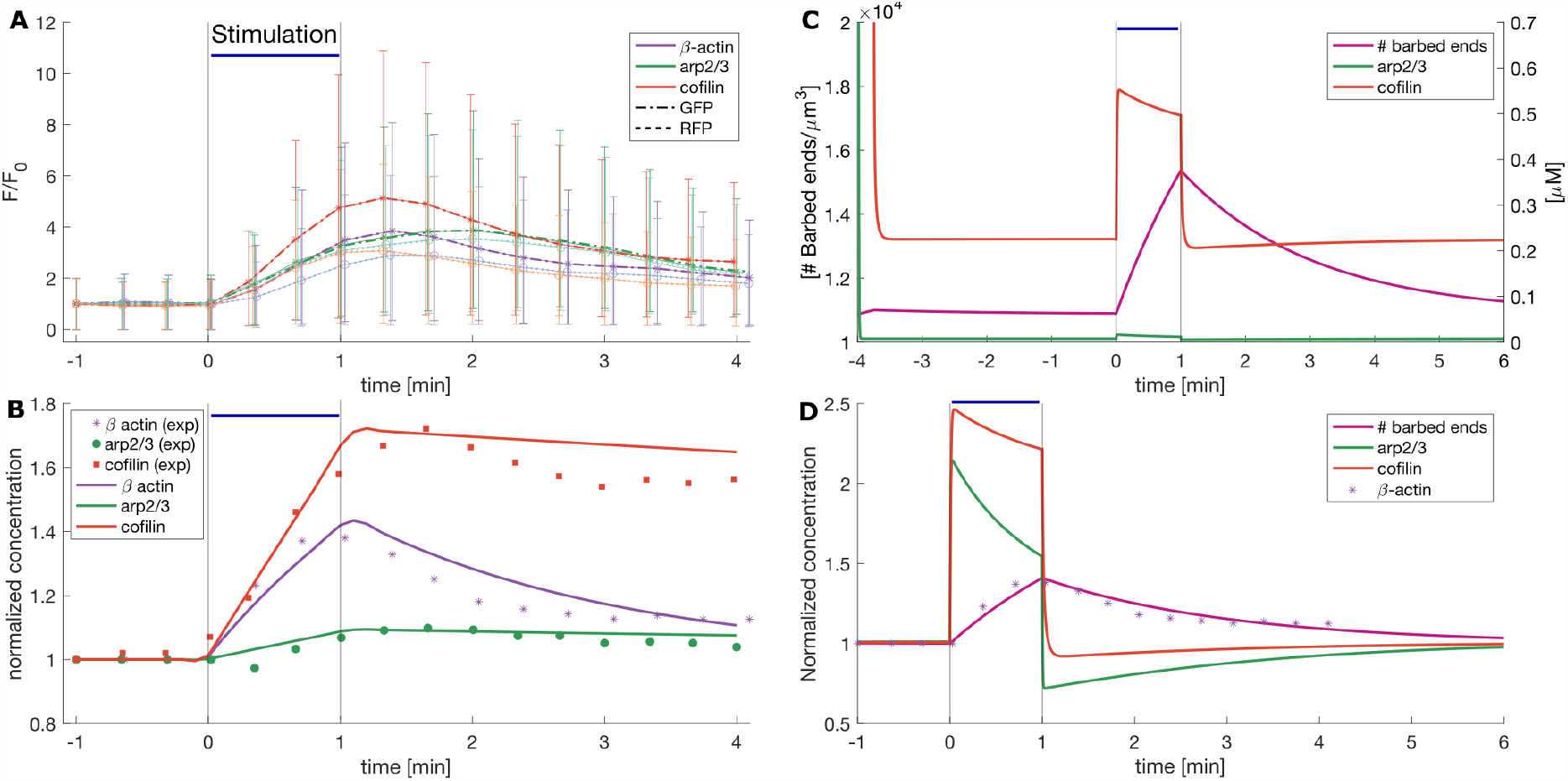
Obtaining model parameters from data. Stimulus is present from *t* = 0 to *t* = 1 minute, indicated by the black vertical lines in the plot. During this time, there is glutamate uncaging and the stimulus-triggered influx *I*_*S*_ *>* 0. **A)** Experimental data taken from (Bosch et al. 2014). Asterisks and dots correspond to the mean of the total fluorescence intensity (F) over the averaged baseline fluorescence intensity (F0) of GFP and RFP, which gives a proxy for the amount of proteins in the spine and its volume, respectively. Errors bars denote SEM. **B)** Markers: Normalized concentrations calculated as GFP/RFP from (A). Lines: evolution of the normalized concentration of proteins in the model used to fit the influx, efflux and decaying rate parameters (squared norm of the residual = 0.0538). **C)** Evolution of the non-spatial version of the model in Eqs. (10), (14), and (15). Note that the left y-axis shows the units of barbed ends per µm^3^ and the right y-axis the units of Arp2/3 and cofilin concentration. **D)** Normalized change in concentration of the quantities on (C) over time obtained by dividing the variable value over its value before the stimulation. Asterisks correspond to the *β*-actin data points in (B).

#### 2.3.1 Protein Dynamics

We obtain the data points from Figure 1 in (Bosch et al. 2014) by annotating the plots in Fiji (Schindelin et al. 2012) with the multi-point tool. This figure shows the volume (RFP) and amount of GFP protein quantified by the relative fluorescence intensity (F) to the average baseline (F0). The points are exported to Matlab (MATLAB 2021) using the `ReadImageJROI.m` function (Muir and Kampa 2015). After scaling the points to the corresponding scale set in Fiji, we calculate the normalized concentration of the proteins as the ratio between the protein fluorescence intensity and the spine volume, as in (Bosch et al. 2014) (Fig. 4A).

For estimation of the parameters corresponding to the actin, cofilin, and Arp2/3 influx into the dendritic spine upon LTP, we develop a minimal model that assumes that these proteins are continuously entering and exiting the spine. Moreover, we consider protein degradation and recent experimental findings showing that some proteins, like *β*-actin, are synthesized locally in the spine (Cajigas et al. 2012; Tiruchinapalli et al. 2003). For simplicity, we have gathered the continuous protein influx and synthesis in a source term *I*, and the efflux and degradation in a decaying rate *k*. Besides these continuous fluctuations, we consider a protein influx triggered by LTP induction *I*_*S*_(*t*), only present during the glutamate uncaging (1 minute). We assume that *I*_*S*_(*t*) are constant values and that during the stimulus window, corresponding to the 1-minute of glutamate uncaging, *I*_*S*_(*t*) *>* 0, and *I*_*S*_(*t*) = 0 otherwise. Hence, in this minimal model, the dynamics for the normalized concentrations of *β*-actin (*β*), Cofilin-1 (*C*), and Arp2/3 (*A*) are given by

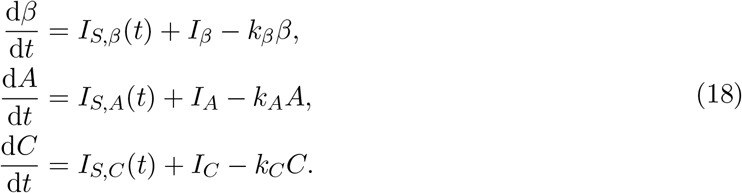

To reduce the number of parameters and guarantee that the pre-stimulus (i.e., *I*_*S,β*_(*t*) = *I*_*S,A*_(*t*) = *I*_*S,C*_(*t*) = 0) level of protein concentration equals one for all the proteins, we first calculate the steady state of the system (*β*^*\**^, *A*^*\**^, *C*^*\**^) = (*I*_*β*_*/k*_*β*_, *I*_*A*_*/k*_*A*_, *I*_*C*_*/k*_*C*_). Then, we scale the system using the nondimensional quantities (*b*(*t*), *a*(*t*), *c*(*t*)) = (*β*(*t*)*/β*^*\**^, *A*(*t*)*/A*^*\**^, *C*(*t*)*/C*^*\**^), which gives

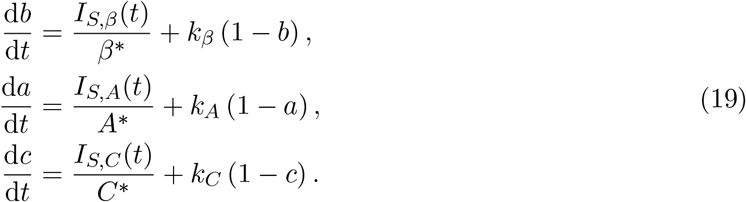

We fit the data points from (Bosch et al. 2014) to the solutions of Eqs. (19) with the lsqcurvefit function in Matlab to obtain the values of the efflux rate constants: *k*_*β*_, *k*_*A*_, and *k*_*C*_; and the influxes upon LTP: *I*_*S,β*_(*t*), *I*_*S,A*_(*t*), and *I*_*S,C*_(*t*). We use the `ode45` solver to evolve the system in Eqs. (19). Figure 4B shows the resulting fit (squared norm of the residual = 0.0538), and the resulting parameters are in Table 1. We obtain the value of actin basal concentrations from (Helm et al. 2021), and for Arp2/3 and cofilin, we take the mean of the molarity of the cytoskeleton proteins, hence (*β*^*\**^, *A*^*\**^, *C*^*\**^) = (3000, 20, 40)µM.

We use a simplified version of our model that neglects the spatial component (i.e., setting to zero the F-actin elongation, repulsive force density, and bulk flow terms in Eqs. 10, 14, and 15) to compare the model output against experimental measurements (Fig. 4C). Note that the evolution of the normalized concentration of barbed ends in the model is similar to the normalized concentration of *β*-actin from the experimental data, despite the differences in the evolution of the concentrations of Arp2/3 and cofilin (Fig. 4D). This discrepancy between the evolution in the experimental data and the model arises because the nucleation and severing events in the model reduce the protein concentrations (terms 2212*f*_*nuc*_ and 2212*f*_*sev*_ in Eqs. (14) and (15), respectively). During our data fitting, we did not account for the interactions between ABPs and actin because the experimental setup does not distinguish when cofilin is bound or unbound to F-actin (Bosch et al. 2014) (Eq. 19). For example, the slow decay of cofilin normalized concentration in the data is attributed to cofilin binding to F-actin and stabilizing it (Bosch et al. 2014). However, in the model, cofilin normalized concentration sharply decays after stimulation to a value below pre-stimulation due to the enhancement of severing events induced by the increase of F-actin. The concentration of Arp2/3 in the model exhibits a similar decay due to the increase of nucleation events promoted by the increase of F-actin. We are interested in the evolution of the spine expansion upon LTP, which we assumed to be regulated by the force generated by actin polymerization. Hence, we considered this model to be a good proxy for the interactions between proteins upon LTP because it exhibits a trend in the evolution of barbed ends similar to the evolution of *β*-actin in the experimental data.

#### 2.3.2 Scale Factors

To convert the concentrations of *β*-actin to number of barbed ends per µm^3^, we assume that there are 167 G-actin per F-actin since the mean length of dynamic F-actin in dendritic spines is around 450 nm (range: 200 - 700 nm, Honkura et al. 2008) and a monomer of actin contributes to 2.7 nm of the filament length (Mogilner and Oster 1996). To change from number of molecules to µM, we use Avogadro’s number and obtain Ψ_0_ ≈ 3.6 number of barbed ends */*(µm^3^µM). To convert number of barbed ends per µm^3^ to concentration of Arp2/3, we follow (Tania, Condeelis, and Edelstein-Keshet 2013) and assume that there is a minimal distance of 37 nm between branches of a filament. Hence, there are 12 molecules of Arp2/3 per F-actin, which gives a scale factor of Ψ_1_ ≈ 0.02µm^3^µM*/* number of barbed ends.

### 2.4 Force-velocity calculation

To calculate the protrusion velocity of the spine (Eqs. 14, and 15), we select a node in the mesh corresponding to the middle of the spine head at the start of the simulation and keep track of its horizontal displacements in a fixed *x*-direction at each time-step. The velocity is calculated by dividing the displacement by the time step duration in every iteration of the model. We assume that negative displacements have zero protrusion velocity. For the force-velocity relationship, the forces are measured locally, i.e., we take the average force generated by the nodes of the triangular face that intersects the cell displacement trajectory.

### 2.5 Stimulus

In our model, the stimulus from the presynaptic terminal triggers an influx of proteins into the spine, consistent with the observations in (Bosch et al. 2014). Therefore, we assume that the stimulus-triggered terms *I*_*S,β*_, *I*_*S,A*_, and *I*_*S,C*_ in Eqs. (10), (14), and (15), respectively, are set to the values in Table 1 divided by the number of basal protein locations during the 1 minute period, and zero otherwise. The division ensures that the total levels of proteins in the model match experimental data. The basal influxes are also divided by the number of basal protein locations. Before and after the stimulus window, these terms are equal to zero. The stimulus-triggered influx is normalized for the spine volume, so the amount of stimulus-triggered influx is independent of the size of the spine. Note that we localize the stimulus-triggered influx to the spine head instead of simulating its transport from the dendrite through the spine neck because the experimental data only shows the increase of the proteins in the spine head. Moreover, the diffusion of G-actin from the shaft to the spine is fast (time constant of 0.005-0.67 s, Honkura et al. 2008), and the available data does not determine whether the proteins enter the spine using vesicular transport or diffusion. Thus, modeling the transport of actin and ABPs through the spine neck would represent an additional delay to the dynamics for which we do not have experimental data. Moreover, most actin filaments in the spine neck are stable and form rings (Bär et al. 2016; Honkura et al. 2008), suggesting that the dynamics between actin and ABPs can differ in that domain.

## 3 Results

Using a minimal model for spine actin-membrane interactions, we investigated the spatio-temporal evolution of the number of barbed ends and concentration of Arp2/3 and cofilin. We used 3D numerical simulations to validate our model against experimental observations of spine growth during sLTP qualitatively. We investigated how different mechanical parameters can affect spine growth dynamics during a single stimulus and finally, predict how spine volume can change due to multiple stimuli. These results are discussed in detail below.

### 3.1 Stimulus-triggered influx reproduces experimentally observed spine growth dynamics

We begin with an investigation of the spatial distribution of the number of barbed ends and the concentration of Arp2/3 and cofilin over time. First, we ran the simulation for 1 minute while keeping the membrane fixed. We observed that the ABPs kept a similar spatial configuration at the end of 1 minute while the barbed ends increased at the center of the spine head (Fig. 5C). We speculate that the increase is due to the repulsive force density term in the barbed ends dynamics (Eq. 10). Therefore, we allowed membrane evolution for 3 minutes (Eq. 3) and investigated whether the barbed end concentration reached a steady state when the force generated by them pushed the membrane forward. We found that the increase in the number of barbed ends slowed down at the end of the 3 minutes. Moreover, the shape of the dendritic spine settled to a new equilibrium shape in which the length of the neck was reduced and its width increased while the width of the head was reduced (Fig. 5A). Therefore, for the initiation of each simulation condition, we use this framework in which mechanical equilibrium is achieved.

**Figure 5:**
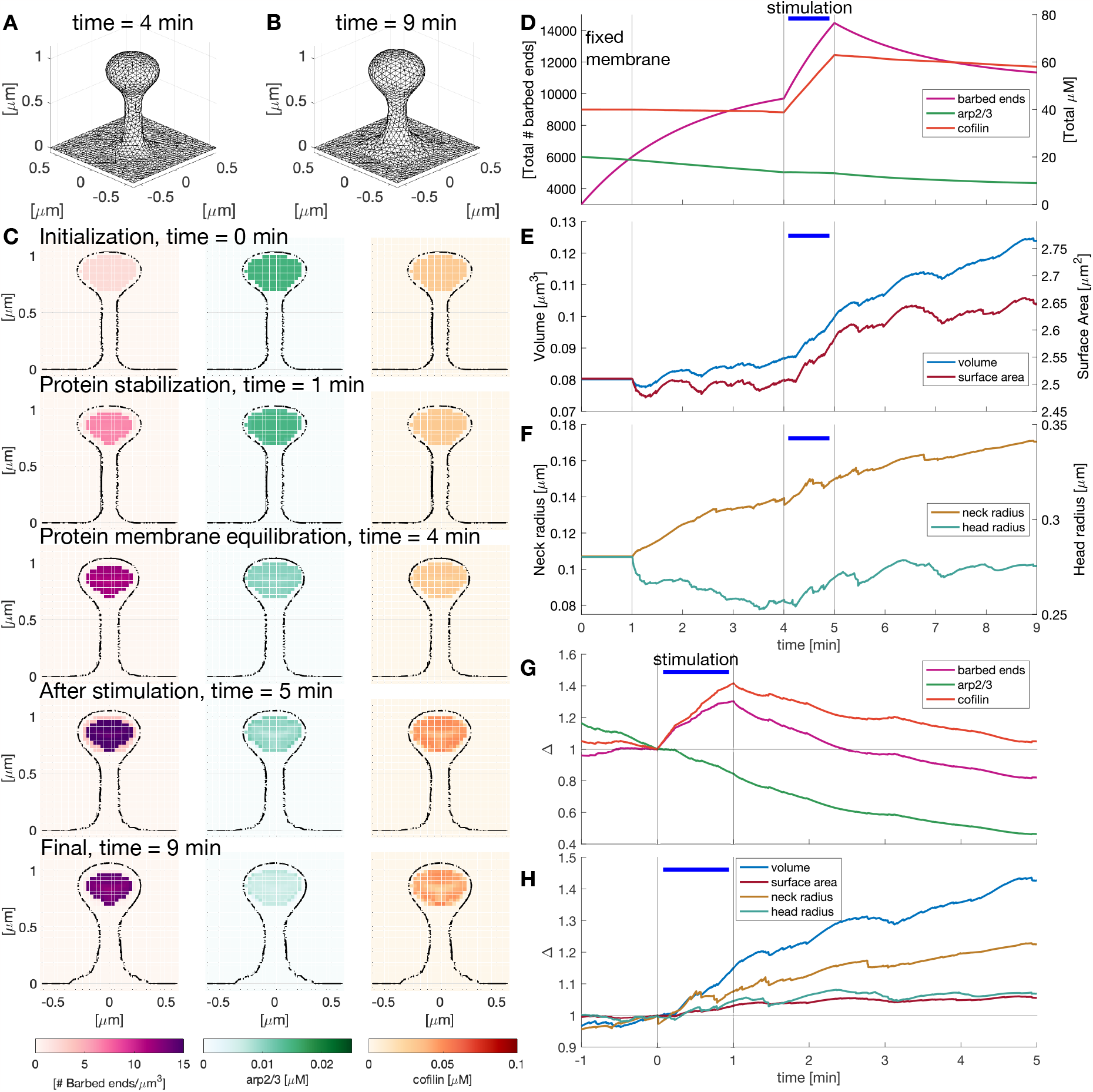
Spatio temporal evolution of proteins upon LTP induction. **A)** Configuration of the dendritic spine before stimulus. **B)** Dendritic spine at the end of the simulation. **C)** Snapshots of the dendritic spine slide at *x* = 0 µm showing the number of barbed ends per unit volume and the Arp2/3 and cofilin concentration at different times. See movie S1. **D)** Sum of the value of the number of barbed ends per unit volume, and Arp2/3 and cofilin concentration in the cubic domain. Vertical black lines signal the times when the simulation changes: at the start, the membrane is fixed, then the membrane is allowed to move until the simulation stabilizes before the stimulus is given (horizontal blue line). **E)** Spine volume and surface area evolution over time. **F)** Spine head and neck radius evolution over time. **G)** Normalized (to *t* = 0 min) change in the number of barbed ends and concentration of Arp2/3 and cofilin in the spine (concentration/volume). **H)** Normalized (to *t* = 0 min) volume, surface area, spine neck, and head radius.

Having established the mechanical equilibrium, we next simulated sLTP induction in the spines by activating the stimulus-triggered influx inside the head (Fig. 2I) for 1 minute, which results in a transient spine enlargement (Fig. 5C). Four minutes after the stimulus, the spines settled to a new larger size with a shorter and wider neck consistent with experimental observations (Bosch et al. 2014; Tønnesen et al. 2014; Yang and Liu 2022). The final shape is shown in Figure 5B.

Next, we integrated the values of the number of barbed ends and concentration of ABPs over time in the cubic domain to understand how the stimulus alters the dynamics of ABPs. Figure 5D shows that the total values of the number of barbed ends, Arp2/3, and cofilin concentration equilibrate before the stimulus is added at 4 min. When the stimulus was triggered, we observed that the number of barbed ends increased, as in the experimental data (Fig. 4A). The cofilin concentration also increased while the Arp2/3 concentration decrease slowed down (Fig. 5D). Arp2/3 decreased throughout the simulation because it is sequestered for nucleation events (see Eqs. 14, 12). After stimulation, the variables decayed at rates similar to the experimental observations (Bosch et al. 2014) (compare with Fig. 4A). These dynamics, informed by the parameter estimation, qualitatively replicate the experimentally observed protein dynamics.

We next calculated the spine volume and surface area over time (Fig. 5E). Note that small fluctuations appeared when we allowed membrane changes (after minute 1) driven by the balance between the force generated by actin polymerization and the force generated by the membrane. The spine volume increased during the stimulus and continued to increase at a slower rate after the stimulus was turned off. The surface area also increased during the stimulus but settled to a new equilibrium value afterward. The first phase of growth is consistent with the main features of sLTP, where the spine head size increases in response to a stimulus (Bosch et al. 2014; Matsuzaki et al. 2004; Tønnesen et al. 2014). However, in experimental data, the spine shrinks after the first phase but we don’t see this shrinkage in our model, which is likely a result of our simplifying assumptions. The addition of further mechanisms to our model could prevent such an increase.

We also measured the radius of the spine neck and head at the same height over time (*z* ∼ 0.63µm and *z* = 0.84 µm, respectively, see Fig. 5F). While the spine head radius showed a similar trend to the spine surface area, the spine neck increased after the stimulus finished, consistent with (Tønnesen et al. 2014). After normalizing the values of volume, surface area, spine neck radius, and head radius (Fig. 5H), we observed an increase in the spine volume (14.92%) and a small increase in the spine surface area (3.03%) and spine head radius (4.6139%) after stimulation. Figure 5G shows the normalized variables of Figure 5D divided over the normalized volume. The increase in cofilin concentration is larger than the increase in the number of barbed ends and similar to experimental data (Fig. 4B). Note that in the model, the decrease of the ABPs and number of barbed ends after stimulation is faster. Moreover, Arp2/3 shows a sustained decrease. Overall, we found that our 3D model qualitatively replicates the temporal trends in protein concentration observed in experiments (Bosch et al. 2014).

To further inspect how the forces generated by the membrane and actin polymerization influence the shape evolution of the spine, we plotted the membrane mesh color-coded by the norm of these forces (Fig. 6A). During the first phase of the simulation, where the membrane is fixed, we observed that the actin polymerization force dominates in the spine head while the membrane force dominates in the spine neck. When the membrane is allowed to move, the actin polymerization force decays at the tip of the spine head, and the membrane force increases at the base of the spine neck. As expected, the force generated by actin polymerization increased in the spine head during the stimulation window but settled to a new steady state after 4 min. Throughout the simulation, the force generated by the membrane is higher at the spine neck and the base of the spine rather than at the spine head. Note that the higher force in the spine neck arises from the smaller radius of the neck compared to the spine head radius while the higher force at the shaft is due to the sharp change in curvature in the junction of the spine neck and the dendrite.

**Figure 6:**
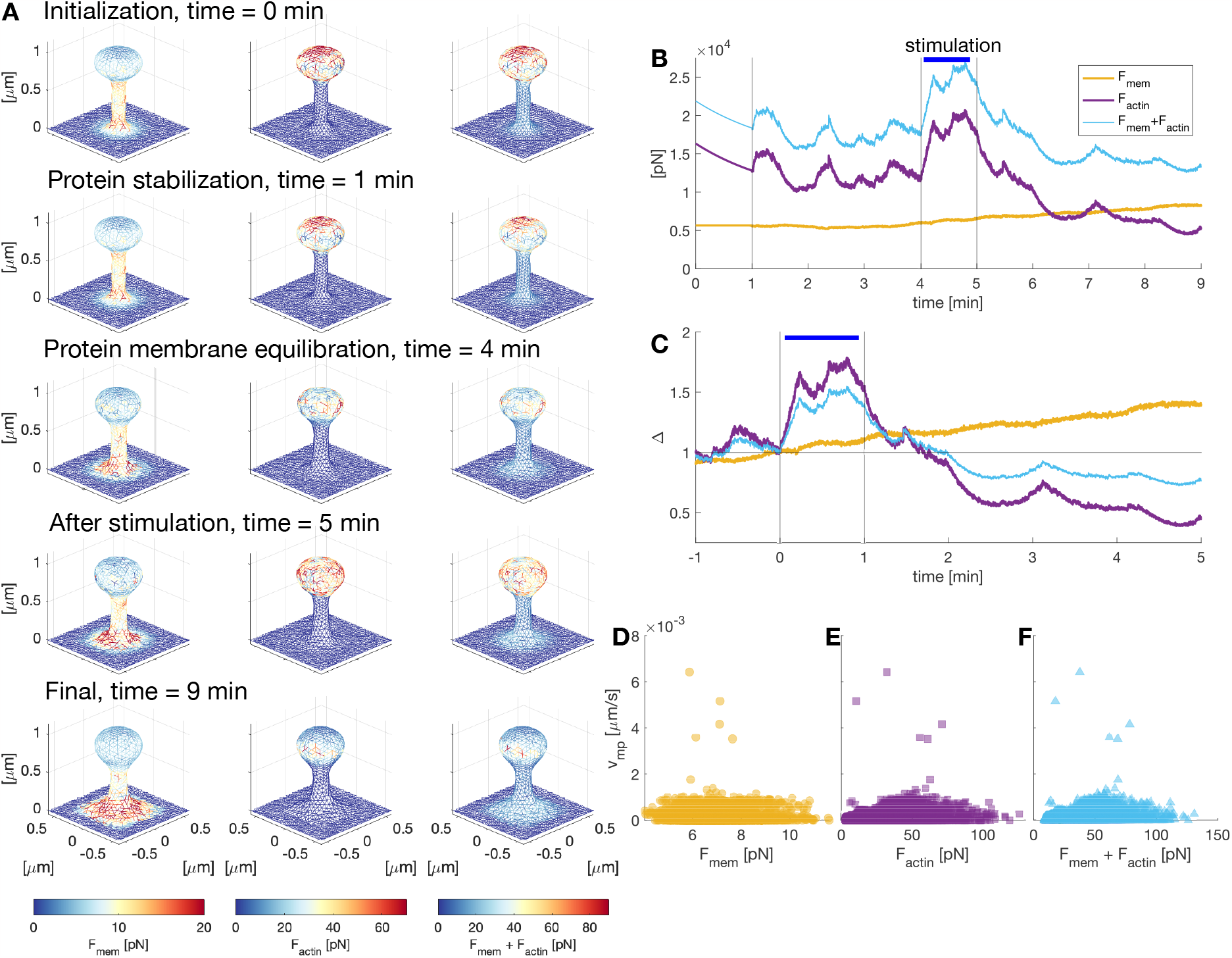
Quantification of spine forces evolution over time. **A)** Temporal evolution of the forces generated by the membrane, actin polymerization, or both, color-coded for the norm of the force vectors. **B)** Sum of the norm of the forces *F* = ||**F**|| generated by the membrane and actin polymerization over the mesh vertices. **C)** Normalized (to *t* = 0 min) total force *F* . **D-F)** Plot of the protrusion velocity over force generated by the membrane, actin polymerization, and both, respectively. The protrusion velocity was calculated as the horizontal displacement of the membrane at the middle of the spine head. The speed with negative directions was set to zero. The force was taken locally, i.e., the force corresponding to the triangular face intersecting the displacement vector.

To quantify the evolution of the forces generated by the membrane and actin polymerization over time, we integrated the norm of the force vectors corresponding to each node of the spine membrane (Fig. 6B). At the start of the simulation, where the spine membrane is fixed, there is a rapid decay of the total force generated by actin polymerization because the barbed ends are pushed back by the repulsive force density (Fig. 5C). The forces settled to a stable value when the membrane was allowed to evolve. During the stimulation window, the force generated by actin polymerization increased while the force generated by the membrane showed a smaller increase. We further analyzed this difference by obtaining the normalized change in the forces (Fig. 6C). Although the total force generated by the membrane has a smaller increase than the force generated by actin polymerization, it decreases the effect of the force generated by actin when the forces are added (Fig. 6C). After stimulation, the force generated by actin polymerization decays to a value smaller than that before stimulation, which relates to the trend shown by the barbed ends (Fig. 5G). The sum of the forces reaches an equilibrium at the end of the simulation.

The force-velocity relationships are shown in Figure 6D-F, for the forces generated by the membrane, actin polymerization, and their sum, respectively. We calculated the force-velocity for spine growth by calculating the membrane horizontal displacement at the middle of the spine head and measured the forces locally. This relationship is nonlinear (could not be fitted to a line). Consistent with other actin-mediated force-velocity relationships, we find that the velocity is higher for smaller forces (McGrath et al. 2003; Brangbour et al. 2011). We observed that higher actin polymerization forces do not correspond to faster protrusions (Fig. 6E), which signals a delay in the membrane response to polymerization forces.

#### 3.1.1 Barbed ends determine spine volume change

The Ca^2+^ entry due to the spine activation of the NMDARs triggers a signaling cascade that leads to activation of Arp2/3 and cofilin (Rangamani, Levy, et al. 2016) (Fig. 1B). Because, in our model, we only account for the external influx of these proteins upon LTP, we investigated whether a further increase due to NMDAR activation enchances spine enlargement. For this, we assumed that the activated proteins contribute to the proteins entering upon LTP induction. Thus, we increased the value of *I*_*S,A*_ and *I*_*S,C*_ by 50%. To have a better representation of the changes in volume, number of barbed ends, and concentration of proteins, we obtained their normalized values at each minute of the simulation (Fig. 7). We found that the increase of Arp2/3 and cofilin (Fig. 7C-D) slightly increases the production of barbed ends (Fig. 7B). However, there is a slight reduction in the normalized volume after stimulation, from 1.1493 to 1.1287 (Fig. 7A). Thus, we concluded that the increase in Arp2/3 and cofilin might produce a negative effect on spine enlargement.

**Figure 7:**
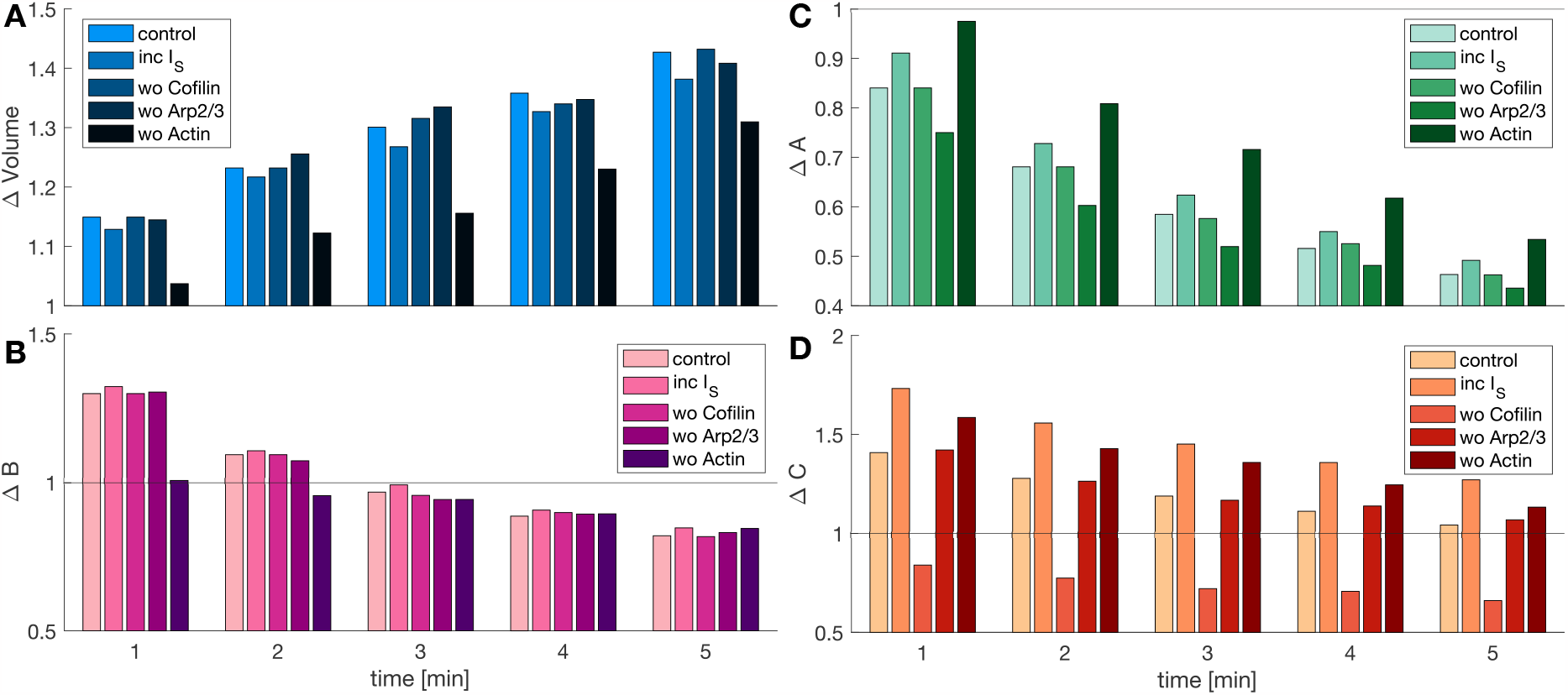
Influence of barbed ends, Arp2/3, and cofilin in spine volume. **A)** Normalized spine volume at different times for various setups where either the stimulus-triggered influx of Arp2/3, cofilin, or actin are excluded, or the stimulus-triggered influx of Arp2/3 and cofilin are enhanced 50%. The stimulus is delivered during the first minute of the simulation. **B-D)** Normalized number of barbed ends per unit volume, Arp2/3, and cofilin concentration at different times.

We then examined the effect of disrupting the stimulus-triggered influx of barbed ends, Arp2/3, and cofilin on spine enlargement. We observed that setting *I*_*S,A*_ = 0 or *I*_*S,C*_ = 0 during stimulation had an insignificant effect on spine enlargement: the normalized increase in volume at the end of the stimulation changes from 1.1493 to 1.1417 and 1.14926, respectively (Fig. 7A). Only impeding the stimulus-triggered influx of actin hindered the increase in spine volume to 3.72% after stimulation. Note that when *I*_*S,β*_ = 0, the peak of normalized Arp2/3 and cofilin concentration after stimulation exhibits an increase, proving that these proteins are sequestered by the stimulus-triggered influx of actin through nucleation and severing events (Fig. 7C-D). From these simulations, we conclude that as long as there are sufficient barbed ends, spine volume change will be robust. The biochemical details of Arp2/3 and cofilin are critical for governing the dynamics of actin reorganization (Pollard and Borisy 2003) but the number of barbed ends determines the growth itself. We inferred that the enhancement of barbed ends after LTP induction is sufficient to overcome the membrane force and allow spine enlargement, in line with previous theoretical work in cell protrusion (Rangamani, Fardin, et al. 2011; Lacayo et al. 2007).

#### 3.1.2 Effect of membrane bending stiffness on spine morphology

The bending stiffness of neuronal membranes varies depending on the neuron type and compartment (i.e., cell body, neurite, growth cone) from 1.8 × 10^*−*19^ to 2.3 × 10^*−*19^ J (Pontes et al. 2013). In our model, we noticed that the total membrane force exhibited only a small increase during the stimulation window (Fig. 6C). However, the total membrane force diminished the increase of the force generated by actin polymerization. Therefore, we further investigated the effect of the force generated by the membrane on spine growth upon stimulation. To do this, we varied the bending stiffness *κ* in Eq. (9) by 25% its value in the simulations. We observed changing membrane stiffness altered the spine shape after stimulation. With a larger membrane stiffness, the spines show a wider neck and thinner head (Fig. 8A), suggesting that the increase in the membrane stiffness counteracts the high curvature of the neck and the side of the spine head. Decreasing the bending stiffness results in spines with a thinner neck (Fig. 8B).

**Figure 8:**
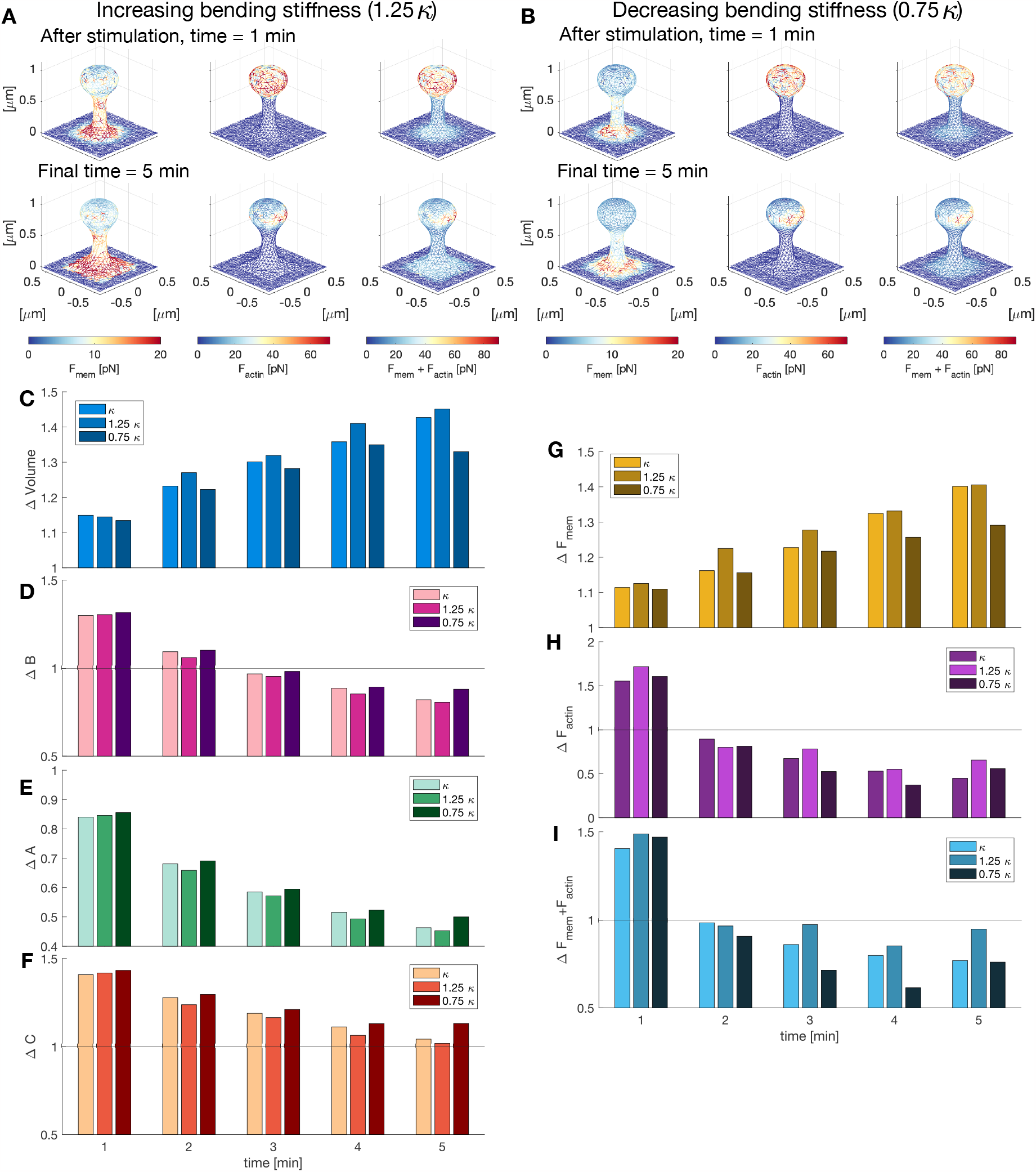
Effect of membrane bending stiffness on dendritic spine expansion. **A)** Dendritic spines after stimulation (top) and at the end of the simulation (bottom), color-coded for the norm of the force vector *F* = ||**F**|| at each node of the membrane mesh. The membrane stiffness *κ* was increased by 25% its value. **B)** Same as (A) but decreasing the value of the membrane stiffness set to 75% its value. **C)** Normalized (to *t* = 0 min) volume for different values of *κ* at different times. **D-F)** Normalized (to *t* = 0 min) number of barbed ends per unit volume, concentration of Arp2/3 and cofilin in the spine for different values of *κ* at different times. **G-I)** Normalized (to *t* = 0 min) total force *F* generated by the membrane, actin polymerization, and both, for different values of *κ* at different times.

Interestingly, we observed that the increase of volume of the spine during stimulation is similar for the different values of membrane stiffness (*V* = 1.1493, 1.1447, and 1.1347 for *κ*, 1.25*κ*, and 0.75*κ*, respectively, Fig. 8C). However, the spine with reduced membrane stiffness showed a smaller change in volume after stimulation due to the reduced spine neck radius. The most notable change in the normalized number of barbed ends, Arp2/3, and cofilin for different values of *κ* occurs after the stimulation window (Figs. 8D-F). The normalized concentrations of *B, A*, and *C* are smaller for increased bending stiffness. There is an increase in the force generated by actin polymerization upon stimulation when the membrane stiffness increases (Fig. 8H). We assumed that such an increase is due to the resistance of the membrane in the middle of the membrane head, where the curvature is smaller and the barbed ends need to produce a higher force to enlarge the spine head. As expected, the changes in the force generated by the membrane are directly related to the membrane stiffness: when the membrane is stiffer it generates a larger force (Fig. 8G). Note that a decrease in *κ* results in a decrease of the overall force (i.e., ||**F**_*mem*_ + **F**_*actin*_||, Fig. 8I). Taken together, we conclude that the resistance offered by the membrane affects the spine shape dynamics and the membrane mechanical properties due to lipid and protein composition could play an important role in sLTP.

### 3.2 Perisynaptic mechanical forces promote spine enlargement

Dendritic spines are embedded in an extracellular matrix and surrounded by other neurons and glia cells (Fig. 9A). Hence, spine enlargement can be promoted or hindered by these perisynaptic elements (Dityatev and Rusakov 2011). Recent experimental results show that coupling F-actin in the spine head with the extracellular space via a molecular clutch is necessary to achieve spine enlargement upon LTP (Kastian et al. 2021) (Fig. 9A). This molecular clutch mechanism was proposed from the observation that the forward movement of the protrusion growth is variable despite the retrograde flow being continuous. Such variation could be to be due to the transient linkage of F-actin with membrane proteins bound to ligands on the substrate (Mitchison and Kirschner 1988). In their experiments, Kastian et al. 2021 showed that shootin1a couples polymerizing F-actin to cell adhesion molecules N-cadherin and L1-CAM. Moreover, LTP induction triggered Pak1-mediated shootin1a phosphorylation, promoting the coupling between F-actin and adhesion molecules. This clutch coupling is thought to reduce the retrograde flow of F-actin and increase the force generated by actin polymerization (Kastian et al. 2021). Here, we investigated the impact of such coupling on volume growth.

**Figure 9:**
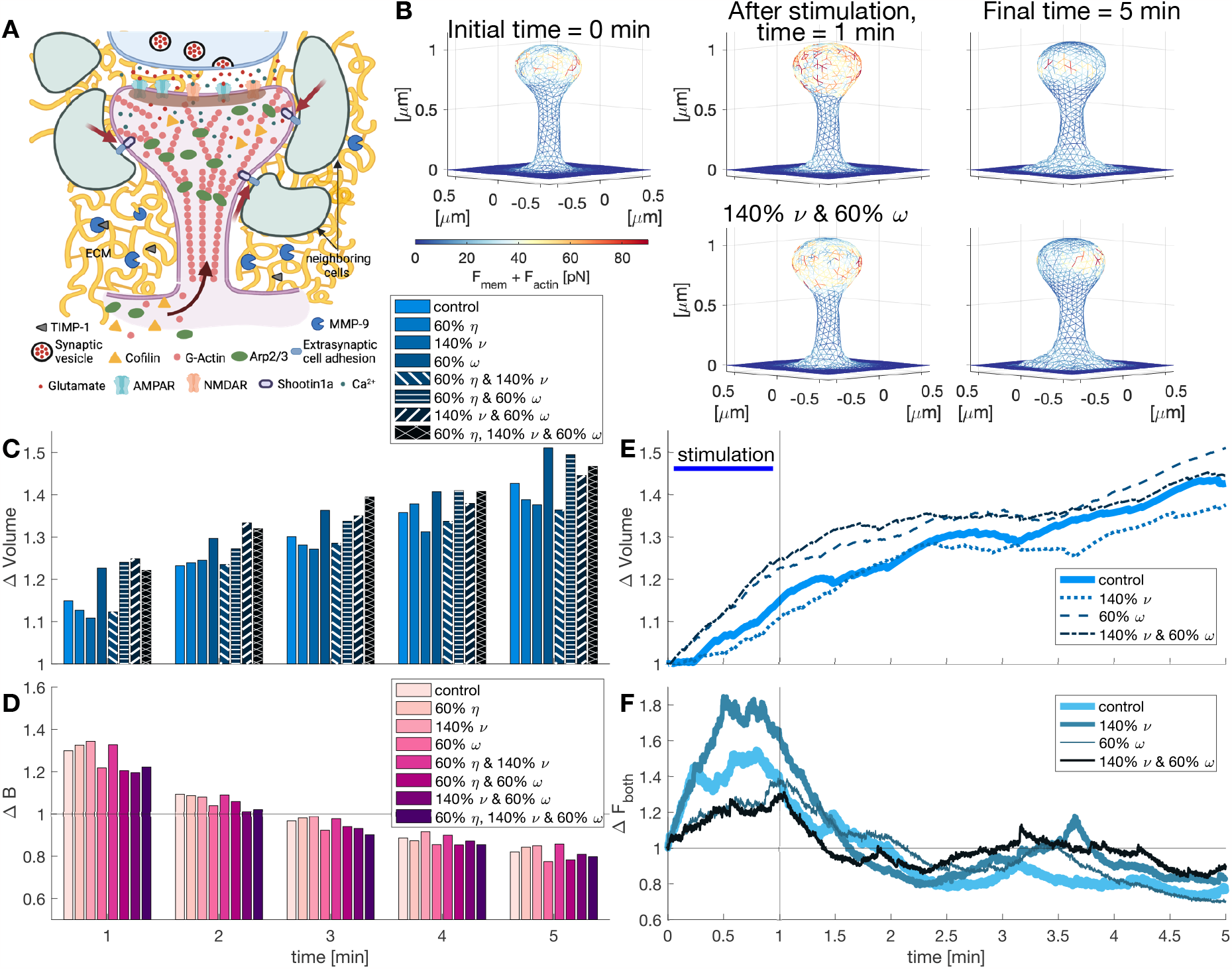
Influence of perisynaptic elements in dendritic spines. **A)** Schematic depiction of coupling between F-actin and ECM and neighboring cells by shootin1a (red arrows). Protease (MMP-9) activity in the ECM. Image created with BioRender.com. **B)** Snapshots of the dendritic spine mesh at different times with control conditions (top) and including the interaction with perisynaptic elements (40% enhancement of the polymerization velocity *ν* and 40% reduction of the drag coefficient *ω*). Edges are color-coded for the combination of the forces generated by the membrane and actin polymerization. For full evolution see Movie S2. **C)** Normalized spine volume evolution at different times. Different colors correspond to different parameter values in the model. **D)** Normalized number of barbed ends per unit volume at different times. **E)** Normalized spine volume evolution over time. **F)** Normalized sum of forces over time *F* = ||**F**_*actin*_ + **F**_*mem*_||.

We assumed that upon LTP induction, shootin1a is highly phosphorylated, and hence, it mechanically couples F-actin to the extracellular adhesive substrates for the extent of the stimulation window. To simulate the reduction of retrograde flow and the resulting enhancement of actin polymerization caused by this coupling, we first decreased the value of the effective filament mobility *η* in Eq. (10). We found that reducing *η* by 40% decreases the normalized volume after stimulation (from 1.1493 to 1.1268) and affects the volume evolution of the spine (Fig. 9C). Therefore, we instead increased the polymerization velocity *ν* by 40% in Eq. (10), which caused a further reduction in the normalized volume at the end of the stimulation window (Δ*V* = 1.1085, Fig. 9C). Since the normalized number of barbed ends per unit volume increases during this period (Fig. 9D), we concluded that the reduced change in volume results from the interaction between forces: when the polymerization force increases, it increases the counteracting forces generated by the membrane. Interestingly, combining the reduction in *η* and the increase in *ν* lessens the volume increment at the end of the simulation (Fig. 9C).

We next investigated which other mechanisms could enhance spine expansion. To allow spine enlargement upon LTP the extracellular matrix remodels through the action of matrix matalloproteinase9 (MMP-9) (Wang et al. 2008). This protease degrades the extracellular matrix after LTP until it is inhibited by the tissue inhibitor of matrix metalloproteinases-1 (TIMP-1) (Magnowska et al. 2016). We mimicked these interactions by decreasing the effective drag coefficient *ω* by 40% during the stimulation window. We observed an enhancement in the normalized volume of 6.73% after the stimulation (Fig. 9C,E). Moreover, when we combined the decrease in the drag coefficient with a rise in polymerization speed, the normalized volume change after the stimulation window was enhanced by 8.65% from control (Fig. 9C,E). Interestingly, the normalized sum of the forces generated by the membrane and actin polymerization reduces (Fig. 9B,F) indicating that the enhancement of spine enlargement after stimulation is achieved by a membrane that is more sensitive to changes in the forces.

### 3.3 Repeated LTP inductions lead to a reduction in spine growth rate

Since synapses receive a series of stimuli (Huang and Kandel 1994), we finally investigated what would happen to spine enlargement upon repeated LTP inductions. To do so, we simulated repeated LTP inductions: the first at the start of the simulation, the second at minute 2, and the third at minute 4. Thus, the spine was stimulated for one minute (i.e., *I*_*S*_ *>* 0 in Eqs. 10, 14, and 15) and let to rest for the following minute. We chose this stimulation protocol because the spine volume increase is reduced after one-minute rest (Fig. 10B).

**Figure 10:**
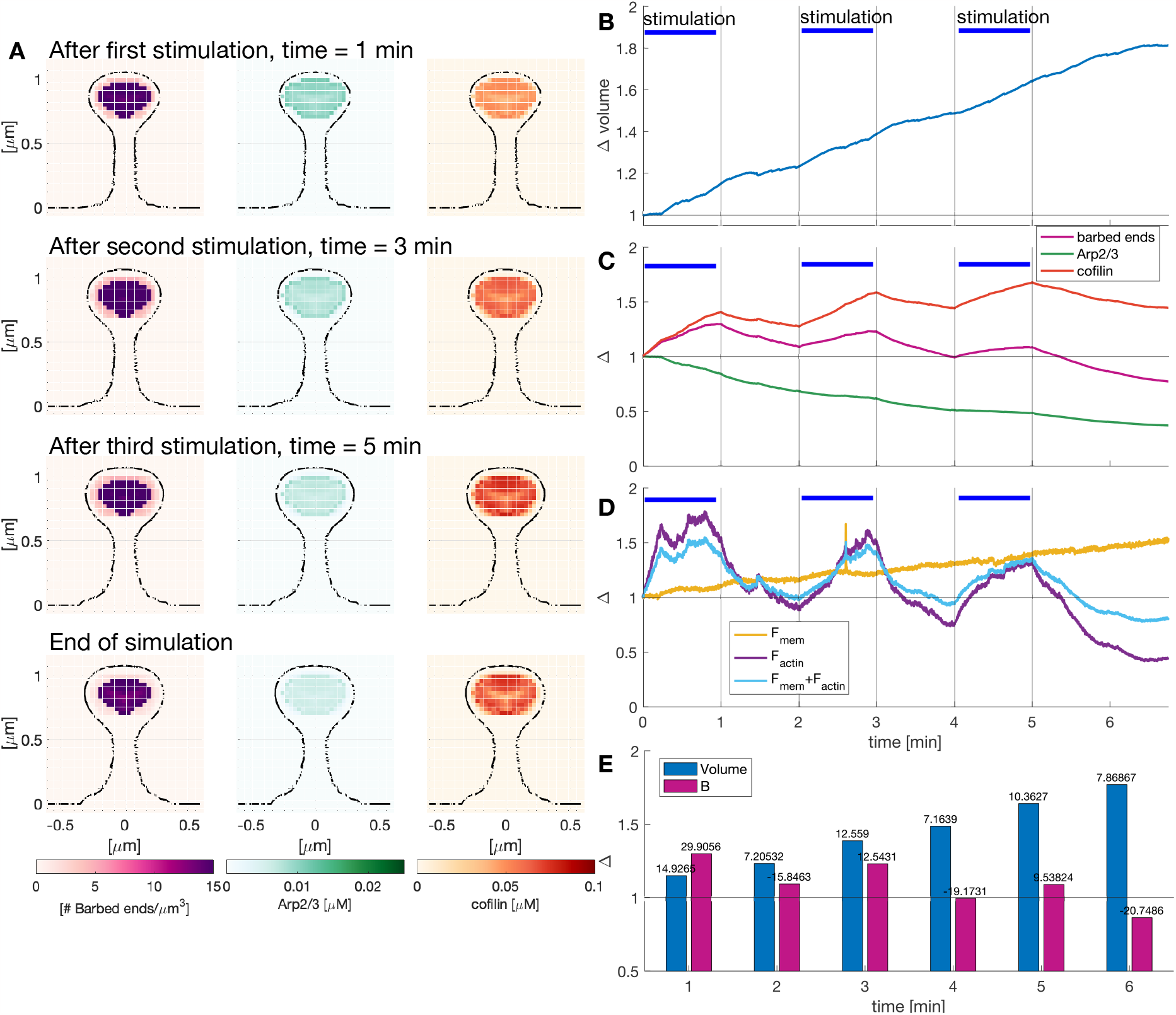
Effect of repeated LTP inductions in protein spatial distribution. **A**) Snapshots of a slide of a dendritic spine at *x* = 0 µm showing the spatio-temporal distribution of the number of barbed ends, Arp2/3, and cofilin. See movie S3 for full evolution. **B**) Normalized volume evolution over time. Different stimulation windows are marked with a blue horizontal line. **C**) Normalized number of barbed ends per unit volume and Arp2/3 and cofilin concentration in the spine. **D**) Temporal evolution of the integral of the force generated by the membrane, actin polymerization, or both, normalized to the value at the start of each stimulus. **E**) Normalized volume and number of barbed ends per unit volume at different times, corresponding to (B) and (C), respectively. Numbers at the top of the bars indicate the percentage change between consecutive times.

We observed an increase in the size of the spine head and a shortening and widening of the spine neck at the end of each stimulus (Fig. 10A). The spine normalized volume increased with every LTP induction (Fig. 10B). Note that the volume increase after each induction is larger than the increase between stimuli (Figs. 10E). However, the volume increase lowers after each instance of the stimuli (14.92%, 12.6%, and 10.36% after the first, second, and third stimulus, respectively). Interestingly, the peak of the normalized number of barbed ends per unit volume was smaller after each stimulus, while the peak in normalized cofilin concentration was higher in the last stimulus (Fig. 10C). The normalized concentration of Arp2/3 reduced its decreasing rate during each stimulation window (Fig. 10C).

The peak of the total normalized actin force during the stimulation window decreases over time while the force generated by the membrane has a steady increase during the simulation that is independent of the repeated stimulations (Fig. 10D). Note that after the third stimulation, the normalized force generated by actin is lower than the normalized force generated by the membrane, which can explain the reduction of the respective volume increase. Taken together, the change in the trends of the normalized protein concentration and forces hint at a complex relationship between the stimulus-triggered influx and the spine volume. In our simulation, spine enlargement was dependent on the spine size at the start of the LTP induction: larger spines showed a smaller increase, consistent with experimental observations (Matsuzaki et al. 2004; Hobbiss, Ramiro-Cortés, and Israely 2018).

## 4 Discussion

In this work, we proposed a minimal biophysical model in which spine enlargement upon LTP is driven by similar mechanisms to those of cell motility. Our model accounts for the spatial localization and chemical reactions of actin barbed ends, Arp2/3, and cofilin, and their interactions with the spine membrane. For purposes of computational tractability, we only focused on a few key ABPs. We chose ABPs that capture minimal actin remodeling events (Pollard and Borisy 2003; Pollard, Blanchoin, and Mullins 2000) and are known to be important for healthy brain function. For example, failure of Arp2/3 function leads to spine loss and abnormal synaptic function, enhancing excitation and leading to similar symptoms to psychiatric disorders (Kim, Rossi, et al. 2015), and experimental studies indicate that cofilin is involved in Alzheimer’s disease synaptic dysfunction (Ben Zablah, Merovitch, and Jia 2020). Importantly, the kinetic parameters of these species were fitted to experimental data (Bosch et al. 2014), indicating that our model predictions represent plausible dynamics. The spatiotemporal maps from the simulations give us a sense of how these molecules may arrange themselves in the spine during active remodeling. It is possible that there may be mislocalization of the proteins in the experiments due to the challenges associated with the overexpression of recombinant proteins fused to GFP (Suratkal, Yen, and Nishiyama 2021), resulting in miscalibration. Nonetheless, the main dynamic events of increase in actin barbed ends and the net force generated are consistent with the literature (Pantaloni, Clainche, and Carlier 2001; Mogilner and Oster 1996).

Our simulations replicate the initial phase of rapid volume increase seen in experiments (Bosch et al. 2014; Matsuzaki et al. 2004; Okamoto et al. 2004). The force-velocity relationships predicted from our model are consistent with other actin-mediated force-velocity relationships (McGrath et al. 2003; Brangbour et al. 2011; Mogilner and Oster 1996). The spatial nature of our model allows us to investigate the forces distribution along the spine membrane over time. We observed that rapid volume increase in our model is driven by the enhancement of the force generated by actin polymerization due to the stimulus-triggered influx of actin. The neck becomes wider and shorter after LTP induction, as observed in experimental data (Tønnesen et al. 2014), hinting that membrane forces could drive changes in the spine neck. Concurrently, the total number of barbed ends decreases below basal levels, thereby reducing the force generated by actin polymerization. Then, the force generated by the membrane dominates, driving spine neck expansion, and hence the slow increase in the total volume of the spine. Therefore, the rapid spine volume increase upon stimulation is due to an enlargement of the spine head by increased actin polymerization while the slow increase in the volume after stimulation is driven by the membrane counteracting the large curvature of the spine neck. Further experiments are needed to test whether the interplay between the polymerization and membrane forces explains the expansion and shrinkage of the spine. Although we did not consider the effect of the periodic actin rings in the neck (Bär et al. 2016; Alimohamadi et al. 2021), we observed that the neck retained its cylindrical structure. Future extensions of our work could explore a previously proposed theoretical hypothesis suggesting that such rings promote mechanical stability of the spine (Alimohamadi et al. 2021).

We showed that the increase of the spine volume upon LTP is enhanced when the interactions between the spine and perisynaptic elements are included. Indeed, recent experimental data found that the clutch molecules, which couple F-actin with the extracellular space, reduce the speed of the retrograde flow and hence, promote the actin polymerization force driving spine enlargement (Kastian et al. 2021). When the coupling with F-actin was disrupted, polymerization of F-actin increased the retrograde flow (Kastian et al. 2021). Furthermore, the activity of MMP-9, which drives extracellular proteolytic remodeling, is necessary and sufficient for spine enlargement and synaptic potentiation (Wang et al. 2008). However, this activity has to be timely inhibited to ensure synaptic responsiveness (Magnowska et al. 2016) hinting to complex dynamics in the ECM.

We examined the spine response under repeated stimuli and observed that their volume expansion reduced after each stimulation. Thus, in line with experimental data (Matsuzaki et al. 2004), larger simulated spines experience less volume increase upon LTP (Matsuzaki et al. 2004). This volume saturation is thought to be regulated by homeostatic mechanisms (Turrigiano 2008), where the spine regulates its synaptic strength by increasing or decreasing the number of AMPARs or NMDARs at the PSD. It has been shown that dendritic spines that experience these homeostatic mechanisms show larger volume increases upon LTP induction (Hobbiss, Ramiro-Cortés, and Israely 2018). Interestingly, spines modulate mechanically their response to multiple stimuli by stiffening (Smith et al. 2007). Here, we show that such homeostasis may be achieved by the interaction between proteins and the forces that drive spine expansion (Fig. 11)

**Figure 11:**
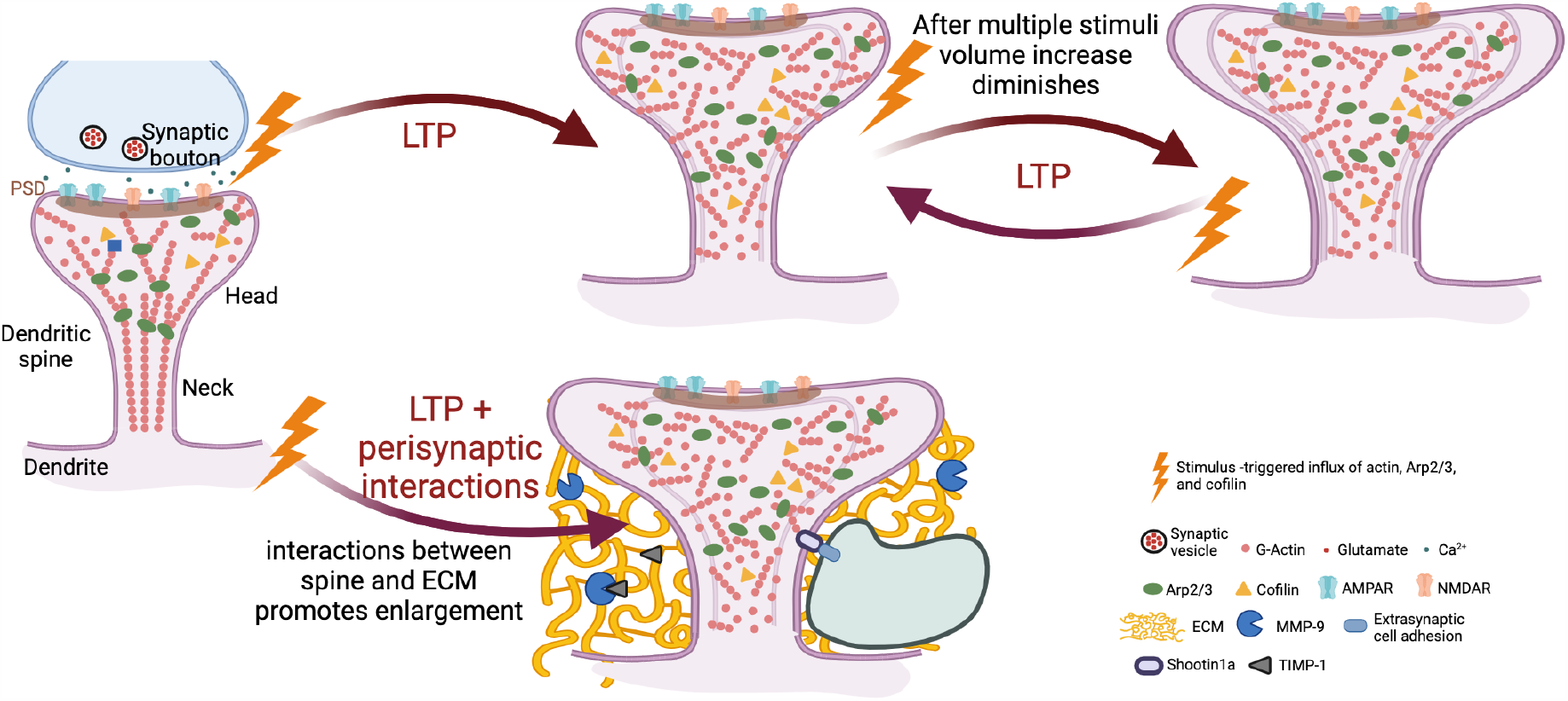
Evolution of the dendritic spine enlargement upon LTP. Summary of the main predictions from our model for the change in spine volume and how it may be regulated by different components. Created with BioRender.com

Computationally, to our knowledge, this is the first 3D model that allows simultaneous protein temporal and spatial evolution, described by partial differential equations (PDEs) in a moving boundary framework, leading to asymmetric shape changes for sLTP. To facilitate computational simulations and mathematical description of the model, we made some simplifying assumptions, such as: 1) There is a large enhancement of protein concentration upon LTP, which allows us to describe their dynamics using a continuum description (PDEs). This limits the model to a short time window after stimulation. 2) Instead of modeling the stimulus-triggered transport of proteins from the dendrite to the spine head through the spine neck, we implemented a localized increment of proteins in the spine head due to the lack of experimental data to determine the type and dynamics of such transport. 3) We assumed that F-actin is constantly branching and severing and only accounted for the number of barbed ends at each location instead of tracking the individual actin filaments. Therefore, our model is not suitable to examine the length or orientation of F-actin. 4) We modeled the dynamics of free cofilin and Arp2/3, and hence, removed the bound cofilin and Arp2/3 through the terms *f*_*sev*_ and *f*_*nuc*_, respectively (see Eqs. 10-15). Although these functions describe the complex binding dynamics between the proteins, our model does not account for the localization of the stable cofilin bound to F-actin. 5) We assumed unconstrained membrane addition through trafficking mechanisms without accounting for the localization of exocytic and endocytic zones (Park et al. 2006), which could influence the resulting spine shape. 6) Instead of modeling the interactions with the ECM and other perisynaptic elements, we modified model parameters. However, molecular clutches are complex and dynamic structures (Giannone, Mège, and Thoumine 2009) and future extensions can incorporate the coupling of F-actin with the substrate through adhesions binding and unbinding, as in cell protrusion models (Sens 2020; Alert et al. 2015). We also focused on the biophysical aspects of spine growth but did not include the signaling events that are a part of the process (Ohadi, Schmitt, et al. 2019; Bell, Bartol, et al. 2019; Bell, Holst, et al. 2022). These signaling events are known to be regulated by the spine shape (Ohadi and Rangamani 2019; Bell, Holst, et al. 2022; Bell, Lee, and Rangamani 2022). In future work, our framework can be extended to include these upstream signaling events and downstream receptor trafficking events. Furthermore, because simulations involved the whole cubic domain, presynaptic and perisynaptic elements can be added at some computational cost. Recent progress in the analysis of moving boundary problems could elucidate an understanding of the relationship between model parameters and its dynamics (Čanić 2021).

In summary, we have shown that the simplest biochemical events associated with actin remodeling result in a volume increase upon LTP, which is enhanced when we account for the interaction with the extracellular matrix. The spine volume increase lessens after multiple stimuli, which hints at a possible homeostatic mechanism by the interaction between the proteins and the forces generated by the membrane. We anticipate that this work will set the stage for coupled modeling and interrogation of the biochemical and mechanical events of sLTP in closer proximity than before.

## Acknowledgements

This work was supported by the NIH Grant Number 1RF1DA055668-01, NIH RO1 GM132106, and by an Air Force Office of Scientific Research Grant FA9550-18-1-0051 to P.R. We thank members of the Rangamani Lab and members of the Le BIP MURI for fruitful discussions. We thank Dr. Christopher Lee, María Hernández Mesa, Sarida Pratuangtham, Dr. Emmet Francis, Dr. Lingxia Qiao, and Yuzhu Chen for their comments.

---

**Movie S1: Protein temporal and spatial evolution upon LTP**. Cross section of the dendritic spine at *x* = 0 µm showing the value of the number of barbed ends (left), Arp2/3 (center), and cofilin (right) concentrations. Volume, surface area evolution, and head radius. Movie corresponding to Figure 5.

**Movie S2: Influence of perisynaptic elements in dendritic spines**. Cross section of the dendritic spine at *x* = 0 µm showing the value of the number of barbed ends (left), Arp2/3 (center), and cofilin (right) concentrations. Volume, surface area evolution, and head radius. Movie corresponding to Figure 9 with increased polymerization velocity 140%*ν* and decreased drag 60%*ω*.

**Movie S3: Effect of repeated LTP inductions in protein spatial distribution**. Cross section of the dendritic spine at *x* = 0 µm showing the value of the number of barbed ends (left), Arp2/3 (center), and cofilin (right) concentrations. Volume, surface area evolution, and head radius. Movie corresponding to Figure 10.

